# *Vibrio cholerae’s* ToxRS Bile Sensing System

**DOI:** 10.1101/2023.05.02.539094

**Authors:** N. Gubensäk, T. Sagmeister, C. Buhlheller, B. D. Geronimo, G. E. Wagner, L. Petrowitsch, M. Gräwert, M. Rotzinger, T. M. I. Berger, J. Schäfer, I. Usón, J. Reidl, P. A. Sánchez-Murcia, K. Zangger, T. Pavkov-Keller

**Author notes:** Nina Gubensäk, Tea Pavkov-Keller. Author Contributions: Conceptualization: T.P.K., N.G., G.E.W., K.Z., J.R. Methodology: T.P.K., N.G., K.Z., T.S., C.B., G.E.W., B.D.G., L.P., M.G., M.R., T.M.I.B., J.S., I.U., P.S.M. Investigation: T.P.K., N.G., K.Z., T.S., C.B., G.E.W., B.D.G., L.P., M.G., M.R., T.M.I.B., J.S., I.U., P.S.M. Visualization: T.P.K., N.G., K.Z., T.S., C.B., G.E.W., B.D.G., L.P., M.G., M.R., T.M.I.B., J.S., I.U., P.S.M. Funding acquisition: T.P.K., N.G., K.Z. Project administration: T.P.K., N.G., K.Z. Supervision: T.P.K., N.G., K.Z. Writing – original draft: N.G., B.D.G., P.S.M. Writing – review & editing: T.P.K., N.G., K.Z., P.S.M., T.S., B.D.G. J.R., M.G., I.U. **Competing Interest Statement:** Authors declare that they have no competing interests. **Classification:** Biological Sciences – Biochemistry.

## Abstract

Cholera represents a diarrheal disease caused by the Gram-negative bacterium *Vibrio cholerae*. Its environmental persistence causing recurring sudden outbreaks is enabled by *V. cholerae’s* rapid adaption to changing environments involving sensory proteins like ToxR and ToxS. Located at the inner membrane, ToxR and ToxS react to environmental stimuli like bile acid, thereby inducing survival strategies e.g. bile resistance and virulence regulation. Currently, transcription factor ToxR is described as main environmental sensor for bile acid, whose activity is enhanced by binding to ToxS. Here, the presented crystal structure of the sensory domains of ToxR and ToxS in combination with multiple bile acid interaction studies, reveals that a bile binding pocket of ToxS is only properly folded upon binding to ToxR. These findings support the previously suggested link between ToxRS and VtrAC-like co-component systems. Besides VtrAC, ToxRS is now the only experimentally determined structure within this recently defined superfamily, further emphasizing its significance.

In-depth analysis of the ToxRS complex reveals its remarkable conservation across various *Vibrio* species, underlining the significance of conserved residues in the ToxS barrel and the more diverse ToxR sensory domain. Unraveling the intricate mechanisms governing ToxRS’s environmental sensing capabilities, provides a promising tool for disruption of this vital interaction, ultimately inhibiting *Vibrio’s* survival and virulence. Our findings hold far-reaching implications for all *Vibrio* strains that rely on the ToxRS system as a shared sensory cornerstone for adapting to their surroundings.

## Introduction

The Gram-negative bacterium *Vibrio cholerae* is the causative agent of the diarrheal cholera disease. Its dangerousness is highlighted by its sudden outbreaks and environmental persistence causing 21 000 to 143 000 deaths worldwide per year (*Ali et al., 2015*) including long-lasting environmental and economic damage (*Kanungo et al., 2022*).

*V. cholerae* exposes a life cycle between dormant and virulent state, enabled by its rapid adaption to changing environments (*Almagro-Moreno et al., 2015b; Bari et al., 2013*). This survival mechanism is maintained via sensory proteins reacting to environmental conditions and substances (*DiRita et al., 1991; Hung and Mekalanos, 2005*). For entero-pathogens like *V. cholerae*, bile acid represents one of the major components for virulence activation, pressuring survival strategies (*Hung et al., 2006; Hung and Mekalanos, 2005; Li et al., 2016; Midgett et al., 2017*).

ToxR is a transmembrane transcription factor (fig. S1) involved in the regulation of numerous genes, not only virulence associated, and can function as an activator, co-activator and repressor (*Bina et al., 2003; Champion et al., 1997; Lee et al., 2000; Morgan et al., 2011; Skorupski and Taylor, 1997; Wang et al., 2002; Welch and Bartlett, 1998*). ToxR periplasmic domain is proposed to act as environmental sensor, being able to bind to bile acids with its periplasmic domain (*Midgett et al., 2017; Midgett et al., 2020*) and consequently activate transcription with its cytoplasmic DNA binding domain (*Gubensäk et al., 2021a; Morgan et al., 2011; Morgan et al., 2019; Pfau and Taylor, 1996; Withey and DiRita, 2006*), thus inducing a switch of outer membrane proteins from OmpT to OmpU (*Simonet et al., 2003; Wibbenmeyer et al., 2002*). Since OmpU is more efficient in excluding bile salts due to its negatively charged pore (*Duret and Delcour, 2006; Simonet et al., 2003*), bile-induced ToxR activation enables *V. cholerae* survival in the human gut (*Wibbenmeyer et al., 2002*).

ToxS is built of a periplasmic, a transmembrane and a short cytoplasmic region (fig. S1). The periplasmic domain of ToxS (ToxSp) interacts with the periplasmic domain of ToxR (ToxRp) forming a stable heterodimer (ToxRSp) (*Gubensäk et al., 2021b*) thereby protecting ToxR from periplasmic proteolysis and enhancing its activity (*Almagro-Moreno et al., 2015a; Almagro-Moreno et al., 2015c; Gubensäk et al., 2021b; Lembke et al., 2018; Pennetzdorfer et al., 2019*).

The exact mechanism of ToxR functionality is not clear yet, so far it is proposed that ToxR binds DNA as a homodimer at the so called ‘tox-boxes’ which represent direct repeat DNA motifs (*Crawford et al., 1998; Goss et al., 2013; Krukonis and DiRita, 2003; Ottemann and Mekalanos, 1996; Pfau and Taylor, 1996; Withey and DiRita, 2006*). The versatile functionality of ToxR suggests a general role of ToxRS in *Vibrio* strains, e.g. the sensing and consequent adaption to changing environmental conditions (*Provenzano et al., 2000*).

Bile also significantly alters the virulence factor production. Nevertheless, the exact mechanism remains unclear. Studies showed opposite outcomes in regard of up- or downregulation of virulence factors by bile (*Bina et al., 2021; Gupta and Chowdhury, 1997; Hung and Mekalanos, 2005; Midgett et al., 2017; Xue et al., 2016; Yang et al., 2013*). On the one hand a bile induced reduction of virulence factor production was observed (*Gupta and Chowdhury, 1997*) via inhibiting ToxT DNA binding ability (*Plecha and Withey, 2015*). On the other hand bile seems to enhance TcpP and ToxR activity (*Yang et al., 2013*) and even induce ToxR ability to activate *ctx* without ToxT (*Hung and Mekalanos, 2005*). Concluding, the effect of bile on *V. cholerae* virulence seems to be complex and probably dependent on multiple other factors e.g. calcium concentration and oxygen levels (*Hay et al., 2017; Sengupta et al., 2014*).

The presented crystal structure of sensory domains of *V. cholerae* ToxR and ToxS reveals ToxS as main environmental sensor in *V. cholerae*. The ToxRS complex exposes a bile binding pocket inside ToxS lipocalin-like barrel that is only properly built via stabilization by newly formed structural elements of ToxR. We performed multiple interaction experiments combined with extensive molecular dynamic MD simulations, to eventually present a bile binding ToxRSp complex, revealing contributions from both proteins to the interaction with bile acid. ToxRSp shows structural and functional similarities with bile sensing complex VtrAC from *V. parahaemolyticus* (*Alnouti, 2009; Li et al., 2016; Tomchick et al., 2016*) thus supporting a common superfamily of co-component signal transduction systems which sensory function is strictly connected to an obligate heterodimer formation (*Kinch et al., 2022*). AlphaFold-Multimer (*Evans et al., 2021; Jumper et al., 2021*) structure predictions furthermore reveal a conserved fold of ToxS in different *Vibrio* species, in contrast to ToxR exhibiting a structural variability among different *Vibrio* strains.

## Results

### Periplasmic domains of *V. cholerae* ToxRS form an obligate dimer

The crystal structure of ToxRp and ToxSp reveals the formation of a heterodimer, with ToxRp contributing secondary structure elements to ToxSp otherwise unstable ß-barrel fold (Fig.1, pdb: 8ALO). Complex formation is achieved spontaneously upon addition of both proteins.

**Fig. 1.**
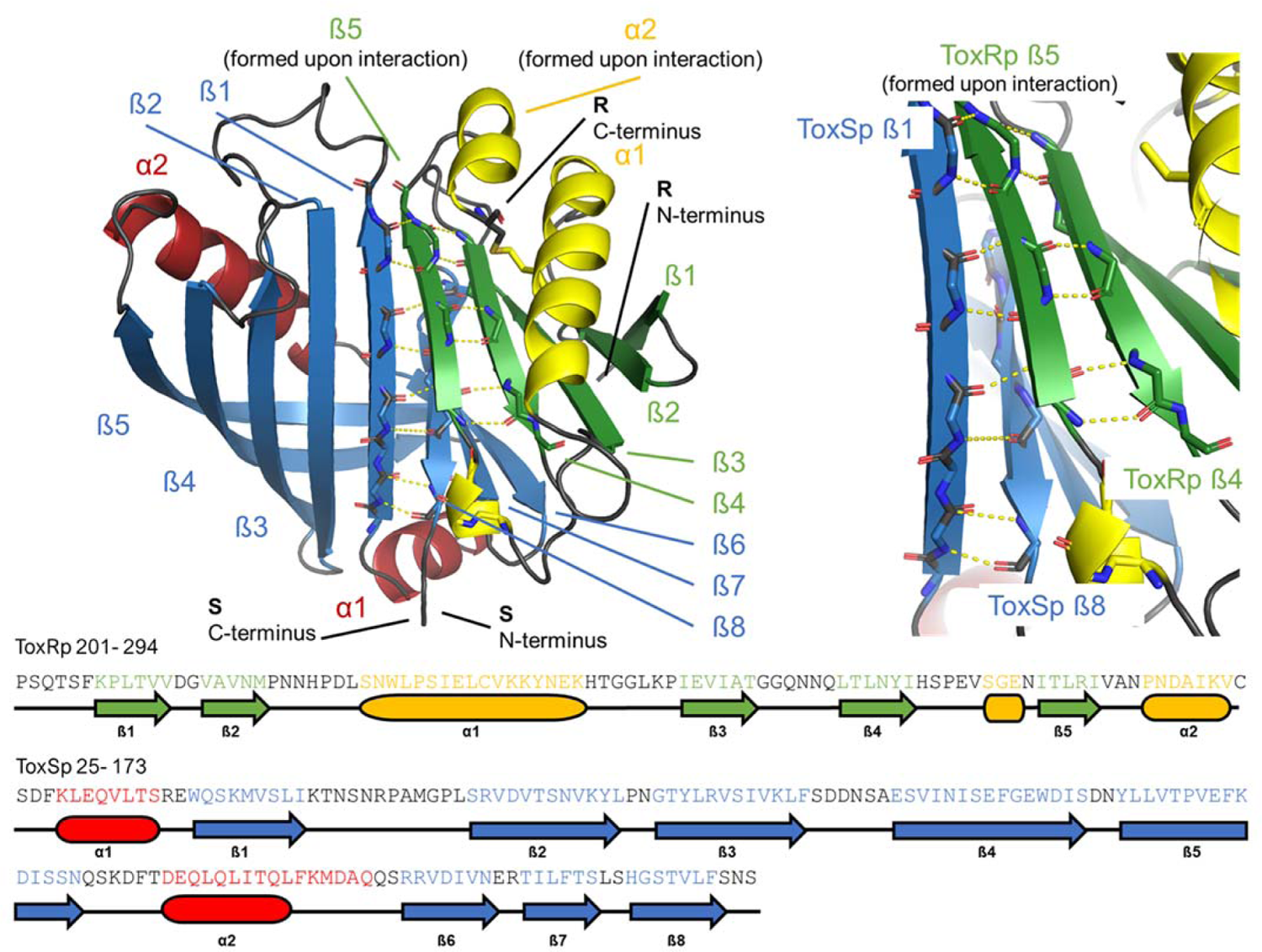
Heterodimer formation of ToxRSp (pdb: 8ALO). ToxRp ß strand 5 completes main chain hydrogen network of ToxSp ß-barrel formation, by interactin with ToxSp ß1. ToxSp ß8 position is stabilized by minor main chain hydrogen bonds with oxSp ß1, as well as side chain interactions with ToxRp ß5 and ß4.

ToxRp in the complex has an αß fold which is formed by an α-helix flanked by an anti-parallel hairpin and a three-stranded anti-parallel ß-sheet, followed by a C-terminal helix. Together, the ß-strands form a five-stranded anti-parallel ß-sheet. The C-terminal Cys293_ToxRp_ is involved in a solution-oriented disulfide bond with Cys236_ToxRp_ of the N-terminal helix.

ToxSp forms a lipocalin-like fold consisting of an eight-stranded ß-barrel stabilized by intermolecular H-bond network, flanked by two α-helices located at the openings of the barrel (fig. S2, S3, table S3). For a correct barrel formation ToxSp ß1 and ß8 would need to be close enough for building main chain interactions. However, ToxSp ß8 contacts only the first three residues of nine residues long ToxSp ß1 (Fig. 1). ToxRp ß5 is positioned into this gap and forms main chain H-bonds with remaining ToxSp ß1 residues. Additionally, close side-chain interactions between ToxRp ß4 and ToxSp ß8 further stabilize ToxSp ß8 positioning (fig. S2, S3). ToxRp ß4 and ß3 are also involved in side-chain interactions with ToxSp ß7 and ß6 completing the strong complex formation. The residue Arg281_ToxRp_ located in ß5 forms a salt bridge with Asp142_ToxSp_ of ToxSp α2, which restricts the opening of the barrel (fig. S2).

There are currently no structures of proteins with significant sequence homology to the ToxRSp complex available in the protein data bank. However, despite of the lack of sequence identity the VtrAC complex of *V. parahaemolyticus* (*Tomchick et al., 2016*) (pdb code: 5KEV, 5KEW) has striking structural and functional similarities with ToxRSp from *V. cholerae* (fig. S18, all-atom RMSD 5.348 Å calculated for 836 atoms; sequence identity: ToxRp-VtrA: 8.1%; ToxSp-VtrC: 11.7%). Similar to ToxRS, the bile sensing functionality of VtrAC induces the type III secretion system 2 (T3SS2), an essential step for virulence activation upon human gastro-intestinal infection.

### Intrinsically disordered ToxRp C-terminus folds into structural elements essential for the complex formation with ToxSp

Small-angle X-ray scattering SAXS experiments indicate a dimer formation of ToxSp in the absence of ToxRp (fig. S4, table S4). Nevertheless, aggregation of ToxSp in solution occurs rapidly suggesting an unstable dimer formation (*Gubensäk et al., 2021b*) (fig S4). The instability of ToxSp can be explained by the inability of the first and the last ß strand of the barrel to form strong hydrogen bonding, thus resulting in an unstable hydrophobic core and consequently aggregation. Interaction with ToxRp likely protects otherwise exposed hydrophobic regions of ToxSp.

Comparison of the ToxRSp crystal structure with our previously determined NMR structure of ToxRp alone (pdb: 7NN6) (*Gubensäk et al., 2021b*), shows the formation of additional secondary structure elements (fig. S5). Unbound ToxRp has a long intrinsically disordered C-terminus in solution, containing the second cysteine Cys293_ToxRp_ forming a disulfide bond with Cys236 _ToxRp_ located in the middle of helix 1. When bound to ToxSp, the unstructured region of the C-terminus of ToxRp performs a disorder to order transition, by forming two additional structural elements: short helix 2 and ß strand 5 (fig. S5). The newly formed short helix 2 contacts the turn between ToxSp ß1 and ß2, as well as residues Leu45_ToxSp_ and Ile46_ToxSp_ located at the C-terminal part of ToxSp ß1 thus increasing the interaction interface and further stabilizing the fold. Although the C-terminal region undergoes extreme structural changes upon complex formation, the orientation of the disulfide bond does not change drastically.

ToxRp ß strand 5 forms the essential basis for the interaction, due to the hydrogen bonding network with ß strands 1 and 8 of ToxSp. ToxRp ß5 also builds a hydrogen network with ToxRp ß4, whose polar regions are otherwise pointing towards the solution, whereas hydrophobic parts contribute to the hydrophobic core. The conformation of ß4 shows only slight changes when interacting with ß5 or its aqueous environment.

### ToxSp in complex with ToxRp contains a bile binding pocket

Bile interaction was tested with ToxRSp using NMR (*Becker et al., 2018*) (saturation transfer difference STD (fig. S6, S7), chemical shift perturbation CSP (fig. S8) and diffusion ordered spectroscopy DOSY (fig. S9)), as well as isothermal titration calorimetry ITC (fig. S10), microfluidic modulation spectroscopy MMS (Fig. 3), size exclusion chromatography with multi-angle static light scattering SEC-MALS (fig. S11) and SAXS (table S4, fig S4). Interaction experiments were performed with bile acid sodium cholate hydrate. All mentioned experiments confirm an interaction of the ToxRSp complex with bile. ITC experiments reveal an intermediate binding affinity of sodium cholate hydrate to ToxRSp (Kd = 9.5 ± 4.4 µM) (fig. S 10).

The lipocalin-like fold of ToxSp forms a hydrophobic cavity (fig. S12), similar to the bile binding pocket of VtrC in complex with VtrA from *V. parahaemolyticus (**Tomchick et al., 2016**).* Using extensive MD simulations enabled a detailed analysis of the protein-ligand interface. Upon binding of bile acid mainly ToxSp loop regions between β1/β2 and α1/β2 undergo major conformational changes (Fig.2) also visible in microfluidic modulation spectroscopy MMS spectra (Fig. 3). The nature of the interactions of bile acid with the protein complex are mostly hydrophobic. The carboxylic moiety forms a strong interaction with Arg52_ToxSp_. Additionally, cholate interacts with residues Ser44_ToxSp_, Ile135_ToxSp,_ Leu138_ToxSp_, Phe139_ToxSp_, Val283_ToxRp_ and Arg281_ToxRp_ (Fig. 2). Both proteins contribute to the bile acid interaction, although ToxSp residues are forming a major part of the interface.

**Fig. 2.**
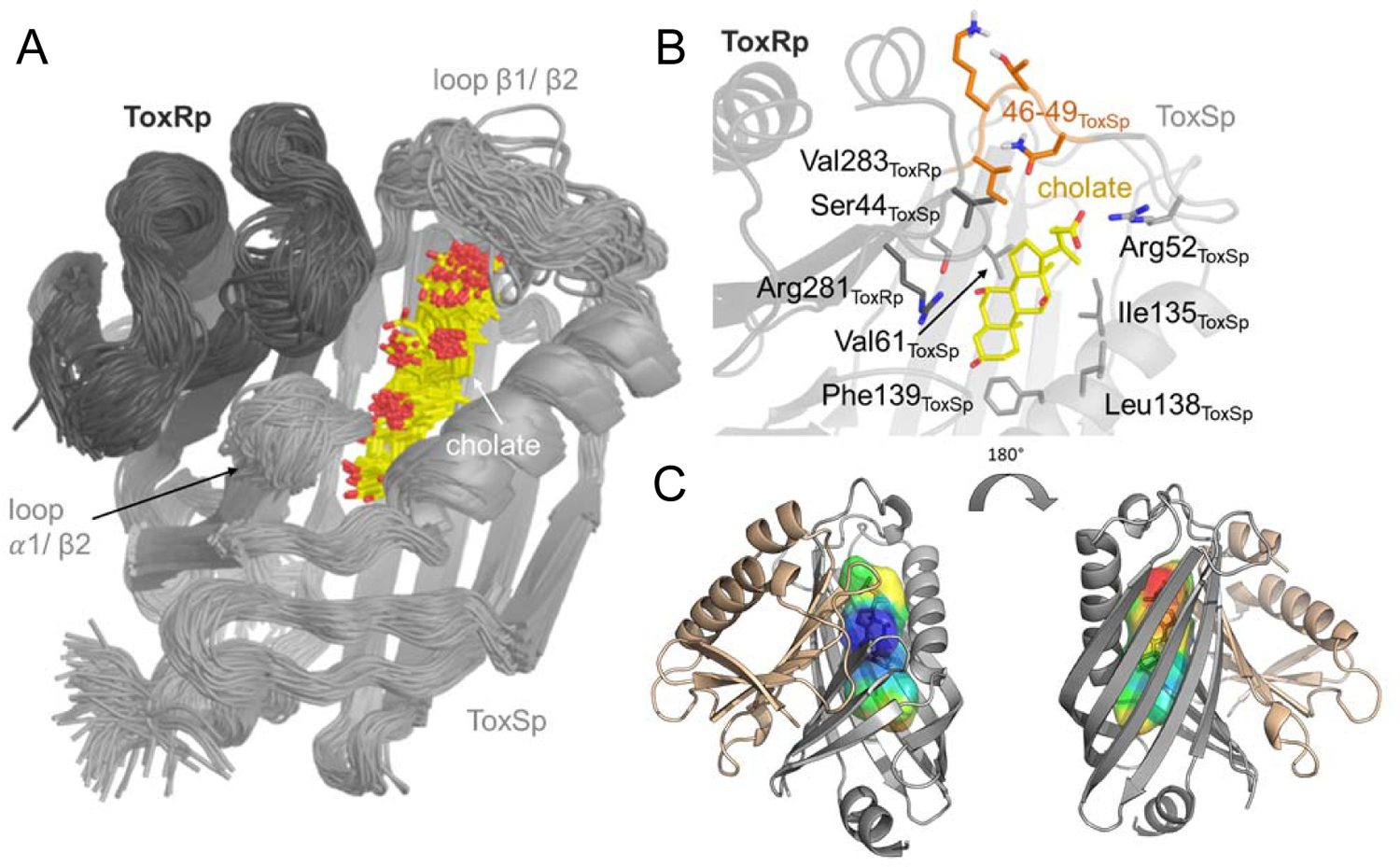
Binding pocket of ToxRSp for bile. **(A)** Superimposition of 100 structures of the ToxRSp complex (dark grey/light grey cartoons) with cholate (sticks, C-atoms in yellow) along one representative classical MD simulation. (**B)** Detailed view of the binding mode of cholate to ToxRSp. The residues with the larger contribution to the binding of the ligand are highlighted as well as residues 46-49 of ToxSp located on loop β1/β2. (**C)** Hydrophobic properties of the bile binding cavity calculated with the CavMan (v. 0.1, 2019, Innophore GmbH plugin in PyMOL). The entrance of the cavity (blue to green) faces the solvent exhibiting a hydrophilic environment, in contrast to the buried hydrophobic areas (yellow to red). For the analysis of the hydrophobicity of the cavities the program VASCo (*Steinkellner et al., 2009*) was used; cavities were calculated using a LIGSITE algorithm (*Hendlich et al., 1997*).

**Fig. 3.**
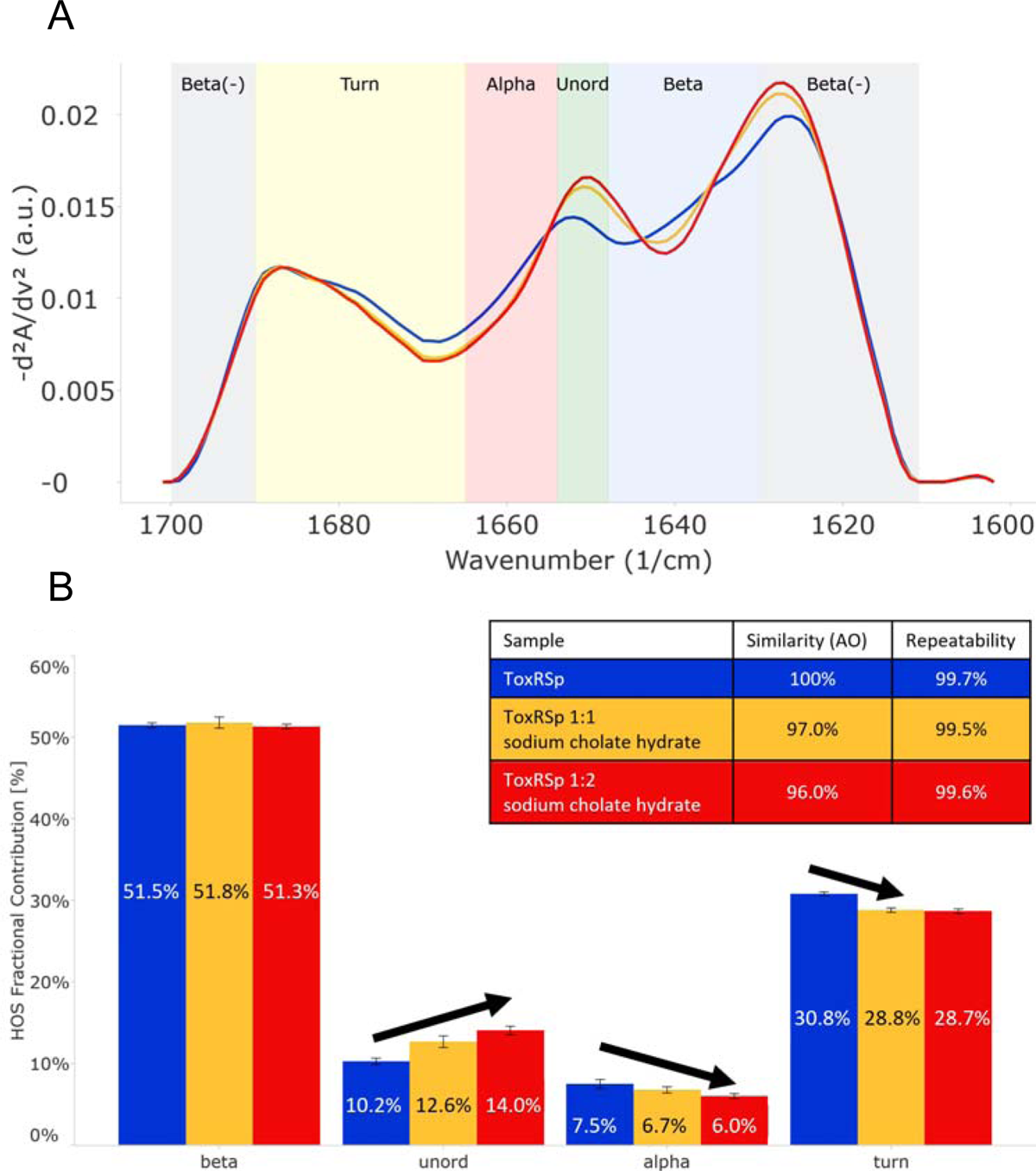
Analysis of microfluidic modulation spectroscopy MMS experiments. MMS experiments of ToxRSp with different sodium cholate hydrate concentrations show conformational changes upon binding of bile to ToxRSp. Additional structural rearrangements could be detected upon higher additions of bile. A: MMS spectra of ToxRSp with and without bile, showing spectral changes upon different bile additions. B: Bar chart of the relative abundance of secondary structural motifs based on Gaussian deconvolution of the corresponding spectra.

### ToxRSp has additional surface exposed bile binding patches

NMR chemical shift perturbation experiments with ToxRSp and bile result in a loss of signals in the 2D experiments probably due to increased relaxation times caused by an increase of molecular weight (fig. S8). Therefore, chemical shift mapping analysis could not be performed. Nevertheless, protein signals could still be observed in 1D proton spectra (fig. S8) thus suggesting an exchange of ligand in solution.

SEC MALS runs propose an increase of molecular weight of ∼10 kDa (fig. S11) of ToxRSp upon bile addition. NMR DOSY experiments of ToxRSp bound to bile acid clearly show a decrease of the diffusion coefficient, which is also linked to an increase of size and weight (fig. S9). Also, SEC-SAXS experiments indicate an increase of molecular weight and radius of gyration of ToxRSp upon addition of equimolar amounts of bile acid (table S4, fig. S4). Via MMS experiments conformational changes could be detected upon bile acid binding but clearly indicate that no severe structural rearrangements occur e.g., aggregation events. Interestingly, MMS experiments also show additional structural changes of ToxRSp upon higher bile acid concentrations (Fig. 3).

The alignment of 1D NMR spectra of ToxRSp before and after the addition of bile acid reveal partial changes of the chemical shifts due to bile interaction (fig. S8), and similar to MMS does not indicate drastically structural changes. Concluding, the presence of bile acid does not seem to induce major structural changes e.g. dimerization events or loss of structure due to aggregation, more likely it leads to local conformational changes near the binding cavity of the ToxSp barrel, similar to VtrAC complex of *V. parahaemolyticus (**Li et al., 2016; Tomchick et al., 2016**)*.

Prompted by these evidences, further MD simulations of the complex ToxRSp were performed with one cholate molecule at the binding cavity in the presence of 14 additional cholate molecules randomly positioned in the solvent (fig. S13). Four solvent exposed regions of the protein, which show a transient binding of cholate molecules could be identified (fig. S13). Three are located on ToxSp, of which two implicate interactions with ToxRp, and one is exclusively located at ToxRp. These results support the experimental finding of an increase of the molecular weight by the binding of more than one cholate molecule.

### ToxRp ß-sheet forms a low-affinity binding region for bile

The MD determined ToxRp binding region to bile is also confirmed experimentally via chemical shift mapping (*Becker et al., 2018; Williamson, 2013*). The determined dissociation constant of 2.6 ± 1.42 mM (Fig. 4, table S5, fig. S14) exhibits a weak binding affinity of ToxRp to bile but physiologically still relevant (*Qin and Gronenborn, 2014; Sukenik et al., 2017*). Concluding, we propose that ToxRp interaction with bile acid is only relevant upon high bile concentrations.

**Fig. 4.**
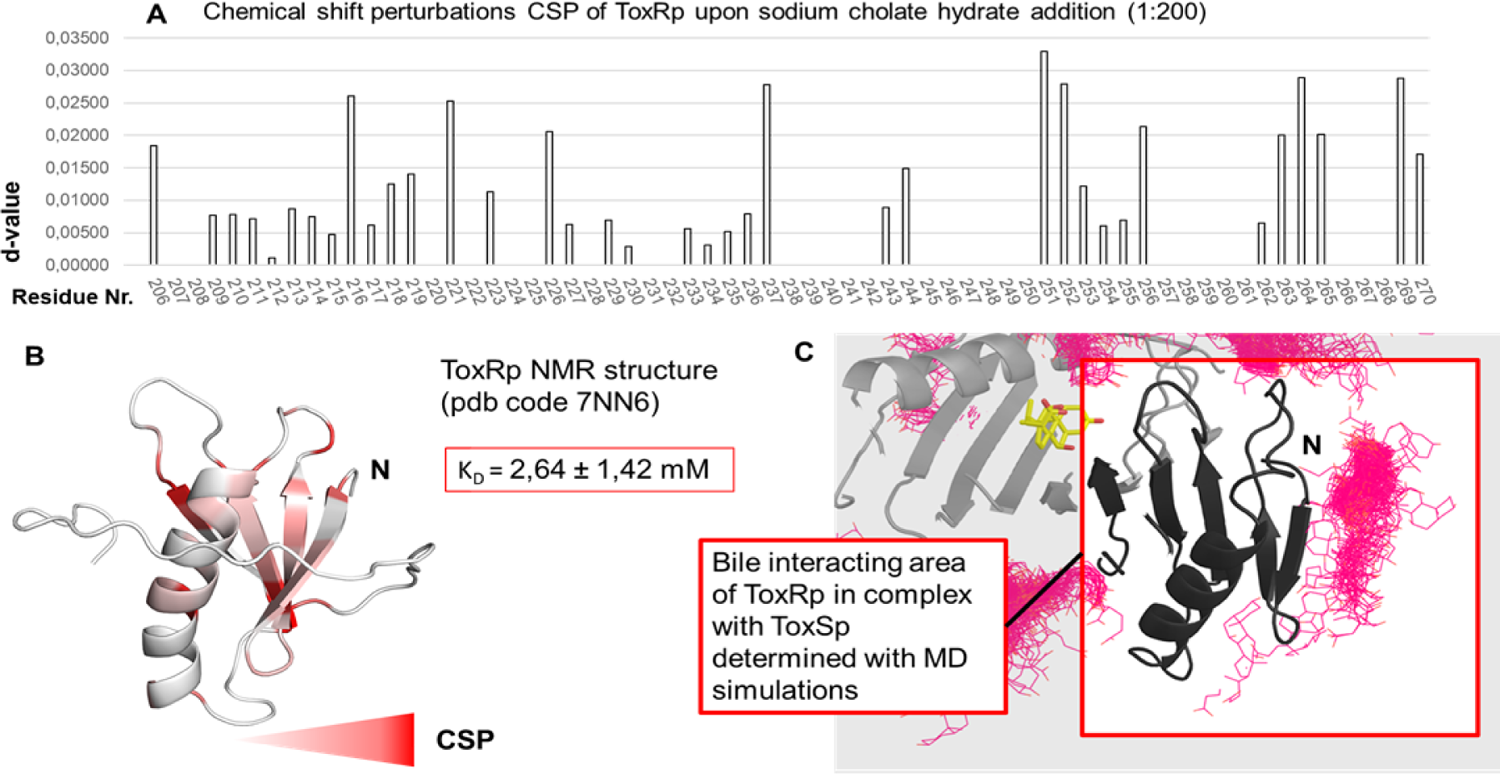
Binding studies with ToxRp and bile. **(A)** Chemical shift perturbation experiments expose residues mostly affected upon binding of sodium cholate hydrate. (**B)** Residues which experience a change of their chemical shift upon bile addition are colored according to a gradient from white to red. Residues highlighted in red are mostly affected upon bile addition and are therefore most likely located at the interaction area. The calculated dissociation constant of 2.6 mM suggests a low affinity binding of bile to ToxRp. (**C)** MD simulations (fig. S13) and CSP experiments propose a bile interacting area located at the ß-sheet of ToxRp.

Interaction experiments with ToxSp alone and sodium cholate hydrate did not reveal any binding event (fig. S7). Due to ToxSp structural instability (*Gubensäk et al., 2021b*) we propose that without stabilization of ToxRp, the binding pocket is not properly formed and therefore binding to bile cannot occur. Additionally, the presented bile binding complex of ToxRSp (Fig. 2) shows that also residues from ToxRp are involved in the bile interaction.

## Discussion

### Bile induced ToxRS activation enables *V. cholerae* bile resistance

Sensory proteins represent an indispensable tool for *V. cholerae* to react to changes in its environment and consequently survive in harsh habitats like the human gut (*Almagro-Moreno et al., 2015b*). The interaction of inner membrane proteins ToxR and ToxS initiates a sensing function followed by signal transmission thereby causing immediate changes in the expression system of the bacterium (*Bina et al., 2003; Childers and Klose, 2007; DiRita et al., 1991*). In order to achieve bile resistance, ToxRS bile induced activation leads to a change of outer membrane protein production from OmpT to OmpU (*Morgan et al., 2011; Provenzano and Klose, 2000; Simonet et al., 2003*).

Our studies reveal a crucial heterodimer formation of the periplasmic sensory domains of *V. cholerae* thereby shaping a bile binding pocket which is only properly folded upon the complex formation. Interaction experiments show that correct heterodimer assembly of ToxRSp is critical for an efficient bile sensing functionality. *In vivo* these findings suggest that bile-induced virulence regulation and OmpU production occurs only when ToxR and ToxS are both present for forming the obligate heterodimer, thus providing a regulatory restriction in the regulation process by bile.

A distinctive feature of members of the ‘ToxR-like’ transcription factor family is the transduction of signals through the membrane without chemical modification, probably via conformational changes (*Eichinger et al., 2011; Gubensäk et al., 2021a; Kenney, 2002; Martínez-Hackert and Stock, 1997a; Martínez-Hackert and Stock, 1997b*). Thus, we suggest that the observed structural changes of the periplasmic domains of ToxRS upon bile recognition are passed on through the ToxR transmembrane domain to its cytoplasmic effector domain, thereby enhancing ToxR binding to recognition sequences and subsequently induction of transcription (Fig. 5).

**Fig. 5.**
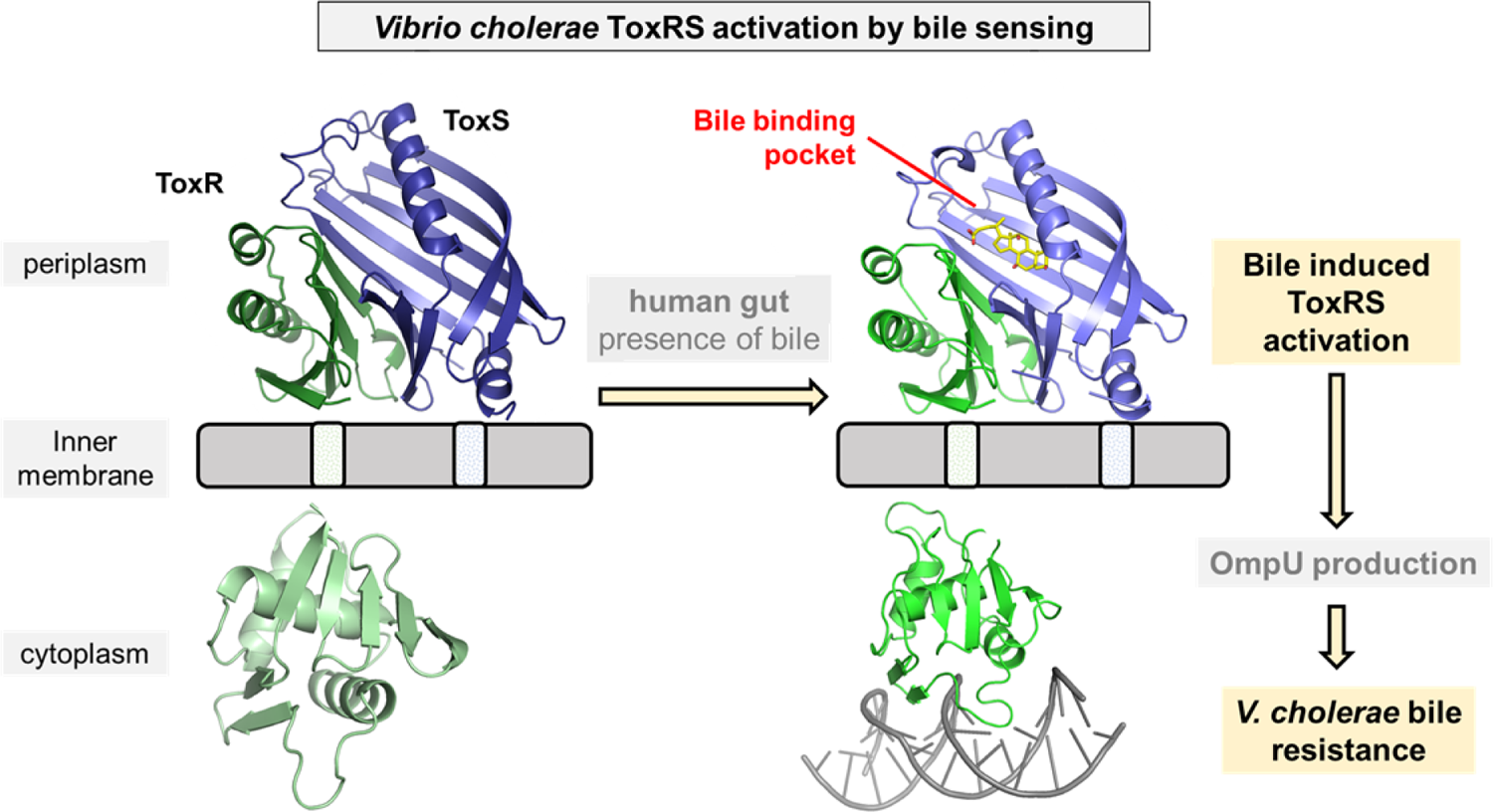
Model of bile induced activation of *V. cholerae* ToxRS. When entering the human gut *V. cholerae* senses the presence of bile acids by binding of bile to the periplasmic domains of inner membrane proteins ToxR and ToxS. The bile binding pocket is formed by ToxS and stabilized by ToxR. Subsequently, the interaction with bile induces ToxR activation which leads to a change of outer-membrane proteins from OmpT to OmpU. OmpU then provides bile resistance and thus enables the bacterium’s survival in the human body.

Higher additions of bile acid induce additional structural changes of ToxRSp as shown in MMS experiments (Fig. 3). A bile acid concentration dependent mechanism could be an efficient tool for *V. cholerae* for sensing its preferred location of infection, the small intestine. In this case only when bile acid concentrations are high enough, the obligatory structural changes could occur, which are transduced through the inner membrane to the cytoplasmic effector domain enabling efficient transcription regulation. Thereby, the high-energy consuming virulent state would only be switched on when *V. cholerae* reaches the small intestine.

The presented experiments do not indicate a bile-induced dimer- or oligomerization of the ToxRS complex, which is currently proposed as active conformation of ToxR. Nevertheless, since experiments are performed with periplasmic domains only, a different behavior of full-length proteins in their natural environment cannot be ruled out. Dimer- and oligomerization events may be dependent on the presence of transmembrane and cytoplasmic domains, and remain to be elucidated.

Taken together we propose that in regard of *V. cholerae’s* virulence expression system ‘ToxR-regulon’ ToxR transcriptional activity is guided by ToxS sensory function. The heterodimer formation is therefore inevitable for the individual functionality of each protein forming a co-dependent system. Understanding the mechanistic details of the ToxRS complex provides a relevant basis for disruption of the crucial interaction interface and consequently inhibition of *Vibrio’s* adaption to its host and its virulent action.

### ToxRS belongs to the superfamily of VtrAC-like co-component signal transduction systems

*Vibrio* pathogens have a complex regulatory mechanism that allows them to survive and cause disease. ToxRS and VtrAC (*Tomchick et al., 2016*) are both *Vibrio* protein complexes fulfilling an essential function since they induce virulence by sensing bile via direct interaction (*Bina et al., 2021; Hay et al., 2017; Hung and Mekalanos, 2005; Tomchick et al., 2016*). Bile sensing is inevitable for the survival of gastro-entero pathogens (*Gunn, 2000; Tomchick et al., 2016*). On one hand outer membrane proteins need to be adapted to bile stress in order to guarantee the survival of the bacteria (*Sistrunk et al., 2016*). On the other hand, the presence of bile signalizes the environment of the host and therefore the virulence activating state needs to be induced in order to effectively colonize the gut.

Structurally, the protein complexes share the ß-barrel lipocalin-like formation of ToxSp/VtrC completed by a ß strand of ToxRp/VtrA, together forming a bile binding pocket (fig. S18). Similar to ToxSp, VtrC is not stable without its interaction partner and tends to aggregate easily (*Tomchick et al., 2016*). There is no structure of unbound VtrA available yet, so we cannot compare if VtrA also forms new structural elements upon binding like ToxRp (pdb code: 7NN6). Another difference between the complexes is that VtrA does not contain cysteine residues like ToxRp. The disulphide linkage represents another option for regulation in *V. cholerae*. Also, bile interaction occurs in a different manner. Whereas in ToxRSp both proteins contribute to the direct interaction with bile, in VtrAC bile binding strictly involves only VtrC (*Li et al., 2016; Tomchick et al., 2016*). If VtrC, in contrast to ToxS, is capable of binding bile acid on its own remains to be elucidated.

Recently, *V. cholerae* ToxRS was proposed as distant homolog to VtrAC, forming together a superfamily of co-component signaling systems which are not linked to sequence homology (*Kinch et al., 2022*). Bacterial two-component systems typically enable signal transduction via chemical modification e.g. phosphorylation of a response regulator (*Mitrophanov and Groisman, 2008*). In the case of VtrAC and ToxRS signal transduction is achieved via binding of signaling molecules like bile acid, thereby inducing conformational changes which likely facilitate transcription regulation (*Eichinger et al., 2011; Gubensäk et al., 2021a; Kenney, 2002; Li et al., 2016; Martínez-Hackert and Stock, 1997a; Martínez-Hackert and Stock, 1997b; Tomchick et al., 2016*).

Although the eight-stranded ß-barrel fold of ToxS and VtrC shows similarities with the calycin superfamily (including fatty acid binding proteins FABP and lipocalins) (table S7) (*Kinch et al., 2022; Tomchick et al., 2016*), the formation of an obligate heterodimer to fold the barrel including the binding pocket is unique for ToxRSp and VtrAC (*Tomchick et al., 2016*).

Despite the obligate heterodimerization and the structural similarities, another feature of this protein family is the arrangement of a two-gene cassette encoding the two proteins (*Kinch et al., 2022*). Using this information, genetic cluster analysis recently revealed additional members of the VtrAC-like superfamily include BqrP/BqrQ from enteric *Burkholderia pseudomallei,* as well as GrvA/FidL from pathogenic *E. coli* O157:H7 saki strain (*Kinch et al., 2022*).

The distinctive structural and functional features of ToxRS and VtrAC support a common superfamily of co-component signaling systems of VtrAC-like protein complexes, whose sensory function is strictly linked to dimerization and ligand binding (*Kinch et al., 2022*). So far VtrAC and ToxRSp remain the only experimentally solved structures of this relatively new defined superfamily.

### ToxRSp obligate dimer formation is conserved among *Vibrio* strains

The predicted protein structures of ToxRSp proteins from different *Vibrio* strains (table S6) share the distinctive ToxRSp fold consisting of a ß-barrel ToxSp and an αß folded ToxRp (Fig. 6A, fig. S15). Whereas ToxRp structure varies among the strains, the barrel shaped lipocalin-like structure of ToxSp is consistent, clearly visible in the alignment of all structures (fig. S15). Sequence alignments (fig. S16 and S17) reveal numerous conserved residues of ToxSp compared to more diverse ToxRp (Fig. 6).

**Fig. 6.**
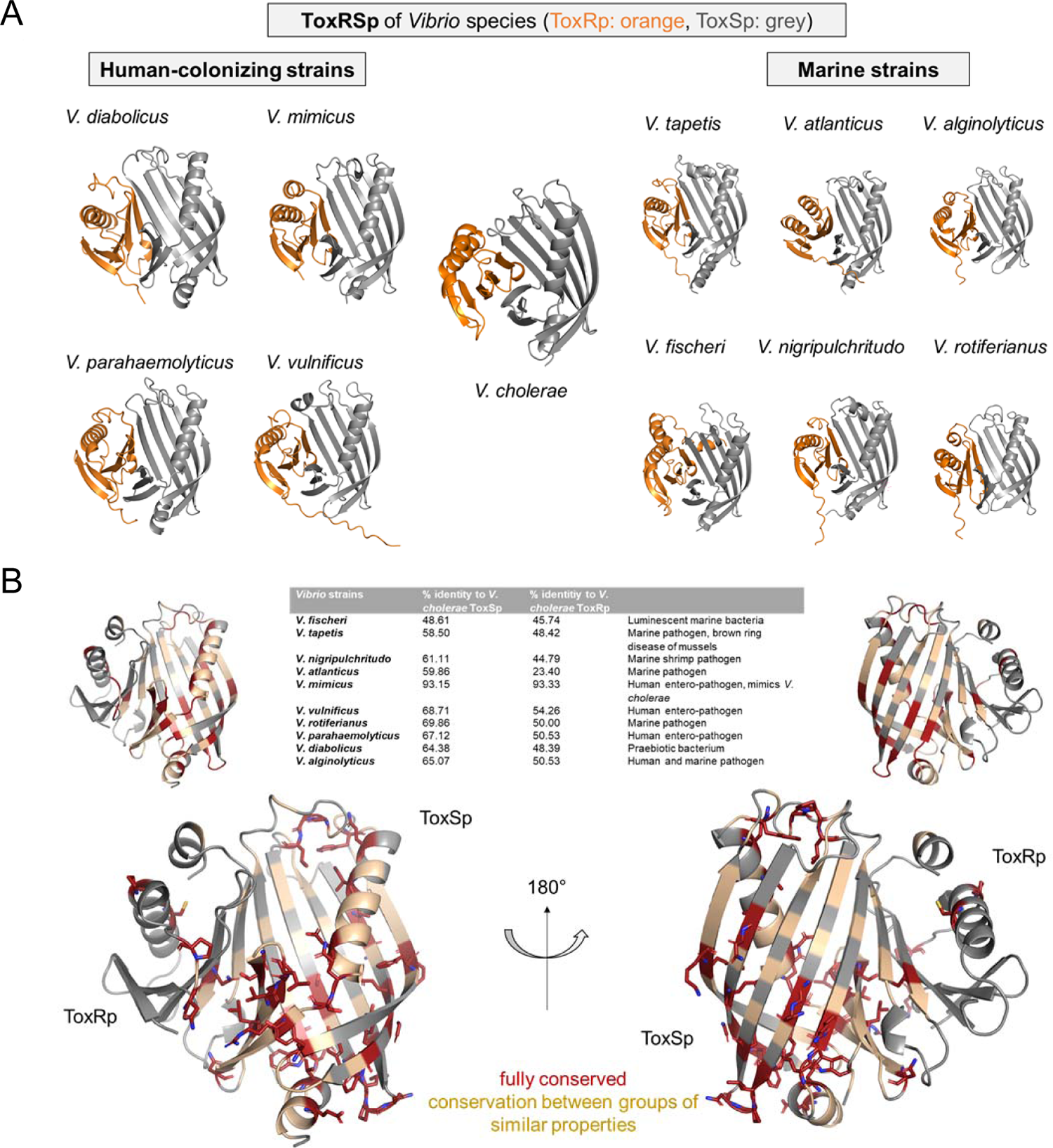
Structural and sequential analysis of sensory domains of ToxRS from different *Vibrio* strains. A: Heterodimer structures of periplasmic sensory domains of ToxRS from *Vibrio* organisms calculated using AlphaFold-Multimer (*Evans et al., 2021; Jumper et al., 2021*). ToxRSp from different *Vibrio* species exhibit similar heterodimer formations. Especially ToxSp ß-barrel structure, which forms the binding cavity, seems to be conserved. **B:** Analysis of conserved residues of ToxRSp. Sequential alignment was done using ToxRS amino acid sequences from Vibrio species (table S6). The sequence identity (%) of ToxR and ToxS proteins from different strains to V. cholerae is shown in the table above the structures. Most conserved regions could be found in ToxSp, especially at the openings of the barrel. ToxRp seems to have a higher sequential variability.

Most of the conserved residues of ToxSp are located at the openings of the barrel as well as ß strands 8, 7 and 6 which are involved in ToxRp interaction (Fig. 6B). Also, residues near the N-terminus, which is probably located near the membrane, seem to be highly conserved in *Vibrio* species. The aspartate residue which is involved in the salt bridge with ToxRp is also one of the conserved residues of ToxSp (fig. S15). The high content of conserved residues in the binding pocket of ToxSp proposes that its general function is to bind to small hydrophobic molecules similar to bile acids.

Compared to ToxSp, ToxRp sequence seems to be only conserved between human colonizing strains (fig. S17). Most of ToxRp conserved residues are involved in the formation of the hydrophobic core, as well as the ToxSp interface (Fig. 6B). Additionally, two cysteine residues are conserved, as well as an arginine/lysine residue involved in the formation of the intermolecular salt bridge with conserved aspartate residue of ToxSp. ToxRp residues involved in the binding to sodium cholate hydrate are also conserved (Val283_ToxRp_, Arg281_ToxRp_).

In conclusion, the sequential and structural alignments reveal that ToxSp is conserved among different *Vibrio* species, whereas ToxRp shows significant sequence similarities only among human colonizing strains. The high number of conserved residues involved in the direct interaction with bile acid furthermore proposes that ToxRSp complex has a common sensory function in *Vibrio*’s.

## Conclusion

Cholera represents a dangerous disease which can lead to sudden epidemic outbreaks due to the high perseverance of its causative *V. cholerae* in aqueous environments (*Ahmed and Nashwan, 2022; Ali et al., 2015*). The bacterium has an immense ability to adapt to challenging conditions due to its sensitive sensory systems. Herein, we present an insight into the regulatory bile binding protein complex of *V. cholerae* sensory proteins ToxR and ToxS. Our analysis reveals ToxS as main conserved environmental sensor in *Vibrio* strains, functionally effective only in complex with transcription factor ToxR. Targeting the disruption of this vital binding mechanism provides a powerful tool for inhibiting *Vibrio’s* adaption to its host and consequently its virulent action.

## Materials and Methods

### Protein expression and purification

All constructs listed in table S1 were generated by using standard procedures and verified via automated sequencing. For protein expression *E. coli* BL21 DE cells were used. Cells were grown under shaking at 180 rpm and induced with 1 mM IPTG when the OD_600_ reached 0.6 to 0.8. ToxRp producing cultures were incubated at 37°C overnight, ToxSp was expressed at 37°C before induction, after growth at 20°C overnight. For production of non-isotopically labeled proteins LB media was used. For isotopically labeled proteins cells were grown in M9 minimal media containing ^15^N labeled (NH_4_)_2_SO_4_ and ^13^C labeled glucose as the sole nitrogen and carbon sources, respectively.

After overnight expression, cell cultures were centrifuged, and the pellet dissolved in 20 ml of loading buffer containing either 8 M urea, 300 mM sodium chloride, 10 mM imidazole pH 8 for ToxRp, or 20 mM Tris, 300 mM sodium chloride, 10 mM imidazole pH 8 for ToxSp. Protease inhibitor mix (Serva, Protease Inhibitor Mix HP) was added to the loading buffer. The cells were disrupted via sonication. The lysate was centrifuged, and the supernatant was loaded on a gravity column containing 2 ml of Ni-NTA agarose beads. The column was washed with 15 column volumes (CV) of loading buffer followed by 5 CV of the loading buffers containing 1 M sodium chloride and 5 CV of the loading buffer containing 20 mM imidazole. His-tagged proteins were eluted with 5 CV elution buffer containing 330 mM imidazole. ToxRp constructs were refolded overnight by dialysis at 4°C in 50 mM sodium phosphate, 300 mM sodium chloride, pH 8. The final purification step includes purification by FPLC using a HiLoad 26/600 Superdex 75 pg column in 50 mM sodium phosphate, 300 mM sodium chloride, pH 8.

### Crystallization & Data Acquisition

Periplasmic protein domains were expressed separately and added in a 1:2 ratio, with excess of ToxSp. Due to ToxSp instability saturation of ToxRp binding sites is only achieved when ToxSp is added in two molar excess. Unbound ToxSp was eliminated by SEC. The protein complex was dialyzed against crystallization buffer (20 mM Tris, 150 mM NaCl, 0.02 % NaN_3_, pH 8) and concentrated to 30 mg/ml using an Amicon Ultra-15 Centricon (Millipore Merck, Darmstadt, Germany). Crystallization experiments were performed with ORYX8 pipetting robot (Douglas Instruments, Hungerford, UK) and sitting drop vapor-diffusion method using 96-well 3-drop plates (SwissCI AG, Neuheim, Switzerland). Initial crystallizations were setup using commercially available screens: Index (Hampton Research, United States), JCSG+ and Morpheus (Molecular Dimensions, United states). Each drop was set up with 0.5 µl protein mix and 0.5 µl of the screen condition. Crystallization plates were incubated at 16° C and H11 condition JCSG+ containing 0.2 M magnesium chloride hexahydrate, 0.1 M Bis-Tris pH 5.5 and 25 % w/v PEG3350 yielded the best diffracting single crystals after two months. The crystals were stored in liquid nitrogen until screening and data collection at the ID30A-3 beamline at the ESRF (Grenoble, France) using a PILATUS detector. Data were collected at 100° K at 0.96770 Å wavelength with 0.20° oscillation range and a total of 700 images.

Data were processed using XDS (*Kabsch, 2010*) and AIMLESS (v.0.7.7.) (*Evans and Murshudov, 2013*). Crystals diffracted up to 3 Å and belong to the space group P6_5_ with cell constants of 72.5 Å, 72.5 Å, 79.4 Å and 90°, 90° and 120°. The structure was solved with molecular replacement using Phaser (2.8.3) (*McCoy et al., 2007*). A partial *ab initio* model of the assembled complex of ToxSp and ToxRp predicted by AlphaFold-Multimer (*Evans et al., 2021*) was used as a template. Employing the complete predicted model of the complex for phasing failed, hence only the larger part of it, ToxSp, was used for initial phasing with Phaser. Furthermore, the predicted model of the ToxSp part was trimmed down at parts below 0.7 predicted IDDT score. The trimmed partial model of ToxSp was suitable to position the model correctly, solve the phase problem and obtain phases that were sufficient enough to place the second molecule, ToxRp, using Phaser. The solved structure was rebuilt and improved in Coot (v.0.9.6.) (*Emsley et al., 2010*) and with ChimeraX (v.1.3.) (*Pettersen et al., 2021*) using the ISOLDE plugin (*Croll, 2018*). Waters were placed manually within Coot. Refinement was performed with REFMAC5 (*Murshudov et al., 2011*). The final refined model was analyzed and validated with PISA (*Krissinel and Henrick, 2007*), MolProbity (*Chen et al., 2010*) and PDB. Data collection and refinement statistics (table S2) were generated within Phenix Table1 (*Liebschner et al., 2019*). The structure was deposited at the PDB with the DOI: https://doi.org/10.2210/pdb8ALO/pdb and the PDB accession code 8ALO.

### Ab initio Models of ToxRp, ToxSp and Dimers with AlphaFold

Ab initio models of ToxRp and ToxSp (*V. cholerae*) were calculated using an AlphaFold 2.1 (*Jumper et al., 2021*) installation with full databases in standard configuration for procaryotes. The search models used for molecular replacement of the heterodimer ToxRSp (*V. cholerae*) as well as the models of the proposed homodimer ToxSp (*V. cholerae*) were calculated on an AlphaFold-Multimer 2.1 (*Evans et al., 2021*) installation with full databases in standard configuration for procaryotes.

For calculation of ToxRSp Heterodimers from various *Vibrio* Species shown in table S6, an AlphaFold-Multimer 2.2 (*31*) installation in standard configuration with full databases and v2 model weights was used. For all species, 5 models were generated and ranked by the highest model confidence metric (0.8·ipTM + 0.2·pTM).

### NMR experiments

All NMR spectra were recorded on a Bruker Avance III 700 MHz spectrometer equipped with a cryogenically cooled 5 mm TCI probe using z-axis gradients at 25° C. All NMR samples were prepared in 90% H_2_O/10 % D_2_O. Spectra were processed with NMRPipe (*Delaglio et al., 1995*).

### Diffusion-Ordered NMR Spectroscopy DOSY

For DOSY experiments all samples were dissolved in the exact same buffer (20 mM Tris, 100 mM NaCl pH 6.5). Sodium cholate was added to ToxRSp samples from a 20mM stock dissolved in the same buffer. The DOSY spectra were recorded using the Bruker dstebpgp3s pulse sequence employing a solvent suppression via presaturation and 3 spoil gradients (*Johnson, 1999; Morris and Johnson, 1992*).

Relevant parameters include a gradient duration of 1.4 ms, a diffusion time of 70 ms and a linear 32-step gradient ramp profile from 1.06 to 51.84 G/cm. Due to phase errors in the first gradient steps, the initial measurement points could not be used, thus the series of points was truncated to afford only the stable signal attenuation. The spectra were processed with Bruker Dynamics Center using standard parameters.

### Saturation Transfer Difference STD

Samples for STD experiments (*Mayer and Meyer, 1999*) contained an excess of ligand (sodium cholate hydrate) compared to protein (protein to ligand ratios: ToxRp 1:50, ToxRSp 1:20, ToxSp 1:20). Three protein selective regions were chosen for saturation: 6000 Hz (amide region), 5000 Hz (amide region) and −5000 Hz (negative control). Experiments with ToxRp and bile were done in two buffers: 20 mM Tris, 100 mM NaCl pH 6.5), and the same buffer that was used for previously published ToxRp NMR interaction studies (*Midgett et al., 2017*) 20 mM KPi pH 8, 200 mM NaCl.

### NMR titration experiments

Titration experiments with ToxRSp were performed in 20 mM Tris, 100 mM NaCl pH 6.5. Titrations with ToxRp and sodium cholate hydrate were carried out in 50 mM Na_2_HPO_4_, 100 mM NaCl pH 6.5. ToxRp experiments were repeated with buffer used by (*Midgett et al., 2017*): 20 mM KPi pH 8, 200 mM NaCl. Sodium cholate was titrated using a 20 mM stock solution prepared in the exact same buffer. 1D ^1^H proton spectra as well as 2D ^1^H-^15^N-HSQC spectra (*Davis et al., 1992*) were recorded for each step of the titrations.

### Isothermal Titration Calorimetry ITC

The isothermal titration calorimetry (ITC) measurements were done using a NanoITC calorimeter (TA Instruments). Proteins and ligand solutions for ITC experiments were prepared in the exact same buffer: 50 mM Na_2_HPO_4_, 300 mM NaCl pH 8.0. The concentration of ToxRSp and ToxRp was 190 µM, sodium cholate hydrate concentration was 1mM and prepared from a 10mM stock solution. All ITC measurements were conducted at 25 °C. In each titration system, the mixing rate was 250 rpm. Injection volume was 2 µL using a titration interval of 20 s. Data were analyzed using the NanoAnalyze software (TA Instruments) and using the One-Site fitting model.

### Size Exclusion Multiple Angle Laser Light Scattering SEC-MALS

SEC MALS experiments were carried out on an ÄKTA pure 25 (Cytavi, Marlborough, USA) using a Superdex 200 Increase 10/300 column (Cytiva) with a flowrate of 0.4 ml/min. The system was connected to a miniDAWN Treos II MALS detector (Wyatt). As sample and running buffer a phosphate buffer (50 mM Na_2_HPO_4_, 300 mM NaCl, pH 8, supplemented with 0.02% NaN_3_) was used. 100 µl of ToxRS without and with bile acid sodium cholate hydrate were measured. Sample 1 was 300 µM of ToxRSp complex, sample 2 was 300µM of ToxRSp with equimolar amount (1:1) of sodium cholate hydrate.

### Microfluidic Modulation Spectroscopy (MMS)

For MMS measurements three 1 ml 5 mg/ml ToxRSp samples were prepared in 50 mM Na_2_HPO_4_, 300 mM NaCl pH 8.0. A 20 mM stock of sodium cholate hydrate was made in the exact same buffer and added to the protein complex for final protein to ligand ratios of 1:1 and 1:2. Additionally, a ToxRSp sample without ligand was measured. For each sample a reference buffer, containing the same amount of sodium cholate hydrate was used.

Samples were measured and analyzed with a RedShiftBio AQS3®pro MMS production system equipped with sweep scan capability (RedShiftBio, Boxborough, MA, US). For background subtraction, chemically identical buffer and buffer-bile mixtures were loaded pairwise with the corresponding sample onto a 96-well plate. The instrument was run at a modulation frequency of 1 Hz and with a microfluidic transmission cell of 23.5 µm optical pathlength. The differential absorbance spectra of the sample against its buffer reference were measured across the amide I band (1,714 – 1,590 cm-1). For each spectrum, triplicate measurements were collected and averaged. The data were analyzed on the RedShiftBio delta analysis software (*Byler and Susi, 1986*).

### SEC-SAXS

SEC-SAXS was performed at the EMBL P12 bioSAXS beamline (PETRA III, Hamburg, Germany). Samples were dissolved in 50 mM Na_2_HPO_4_, 300 mM NaCl pH 8.0 and 3% glycerol to prevent radiation damage. For the single proteins, ToxSp, ToxRp and ToxRSp, a Superdex 75 Increase 10/300 column was used. For the ToxRS complex in the presence of bile acid, a Superdex 75 Increase 5/150 column was used. The column outlet was connected the flow-through SAXS cell.

Throughout the elution, frames (I(q) vs q, where q = 4πsinθ/λ, and 2θ is the scattering angle) were collected successively. The data were normalized to the intensity of the transmitted beam and radially averaged; the scattering of the solvent was subtracted from the sample frames using frames corresponding to the scattering of the solvent. For optimized selection of buffer and sample frames the program CHROMIXS was employed. Data were further analyzed and prepared via PRIMUS from the ATSAS software (*Manalastas-Cantos et al., 2021*), model preparation was done using DAMMIF at the online ATSAS server provided by EMBL Hamburg (https://www.embl-hamburg.de/biosaxs/atsas-online/). Comparison to theoretical scattering curves was performed with CRYSOL. Figures were done using PyMOL (*The PyMOL Molecular Graphics System, Version 1.2r3pre, Schrödinger, LLC.*). The data have been deposited in the SASBDB databank. Accession codes: *tbd*.

### MD simulations

The Cartesian coordinates of the complex between ToxRp and ToxSp were used as initial structure for our MD simulations. The minimum energy structure of the ligand sodium cholate hydrate was first of all minimized with the GFN2-xTB (*Bannwarth et al., 2019*) using CREST, (*Grimme, 2019*), minimized with r^2^SCAN (*Furness et al., 2020*) and the RESP point charges for all atoms of cholate were derived from the electrostatic potential computed at the HF-6-31G* level of theory with Gaussian16 v. C.01 (*Frisch et al., 2016*). The protonation states of the titritable residues on the protein complex were calculated via the H^++^ web server (; *Anandakrishnan et al., 2012; Gordon et al., 2005; Myers et al., 2006*) assuming a pH value of 8.0. Cholate was manually docked at the hydrophobic inner chamber of the ToxRSp complex guided by the superimposition of the crystallographic structure of *Vibrio parahaemolyticus* VtrA/VtrC complex in the presence of taurodeoxycholate (PDB id. 5KEW) (*Alnouti, 2009; Li et al., 2016*).

MD simulations were carried out using the suite of programs AMBER20 (Amber20*, 2022*) Protein residues and solvated ions were treated with the AMBER ff19SB force-field (*Ponder and Case, 2003*) and cholate was described as AMBER atoms types following the standard procedure in antechamber. We simulated two systems: the apo complex (ToxRSp) and the cholate-bound complex (ToxRSp + cholate). The two systems were then energetically minimized to avoid close contacts, and then placed in the center of a cubic box filled with OPC water molecules (*Izadi et al., 2014*). Then, the solvated systems were minimized in three consecutive steps (all protons, solvent and all system) and heated up in 50 ps to from 100 k to 300 K in a NVT ensemble using the Langevin thermostat (gamma friction coefficient of 1.0). Care was taken to constraint the solute during the heating step by imposition of a harmonic force on each atom of the solute of 40 kcal mol^-1^ Å^-2^. Afterwards, these harmonic constraints were gradually reduced up to a value of 10 kcal mol^-1^ Å^-2^ in 4 simulation stages (NVT, 300 K). Then, the systems were switched to constant pressure (NPT scheme, 300 K) and the imposed constraints during the heating step were totally removed. Finally, each of the systems was submitted to three independent MD simulations of 0.8 μs (total simulation time per system: 2.4 μs). Atom-pair distance cutoffs were applied at 10.0 Å to compute the van der Waals interactions and long-range electrostatics by means of Particle-Mesh Ewald (PME) method.

SHAKE algorithm was applied to restrain the hydrogen atoms on water molecules. MD trajectory analysis was carried out using the CPPTRAJ (*Roe and Cheatham, 2013*) module from AMBER20 for monitoring the root-mean-square distance (RMSD) and root-mean-square fluctuation (RMSF), amongst other parameters.

## Acknowledgments

We are grateful for the beam time on ID30A-3 at ESRF (Grenoble, France) for intensive diffraction screening and data collection, as well as for staff support during data collection measurements (BAG mx-1740). J.R., K.Z. and T.P.K. acknowledge the support of the field of excellence BioHealth at the University of Graz.

## Supporting Information for

**Fig. S1.**
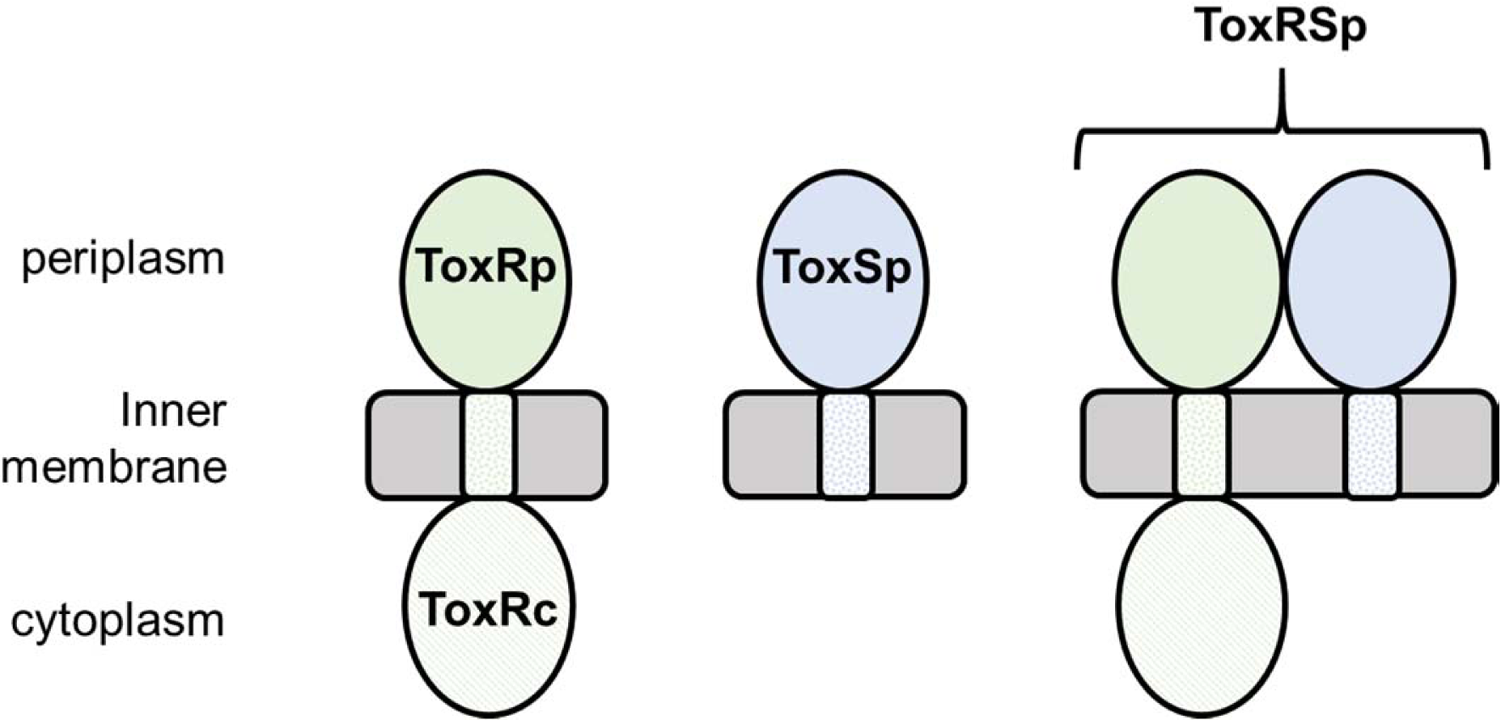
Inner membrane spanning domains of *Vibrio cholerae* regulators ToxR and ToxS. ToxRp, ToxSp stand for the periplasmic domains of ToxR and ToxS, respectively. ToxRSp is the complex formation of periplasmic domains of ToxR and ToxS.

**Fig. S2.**
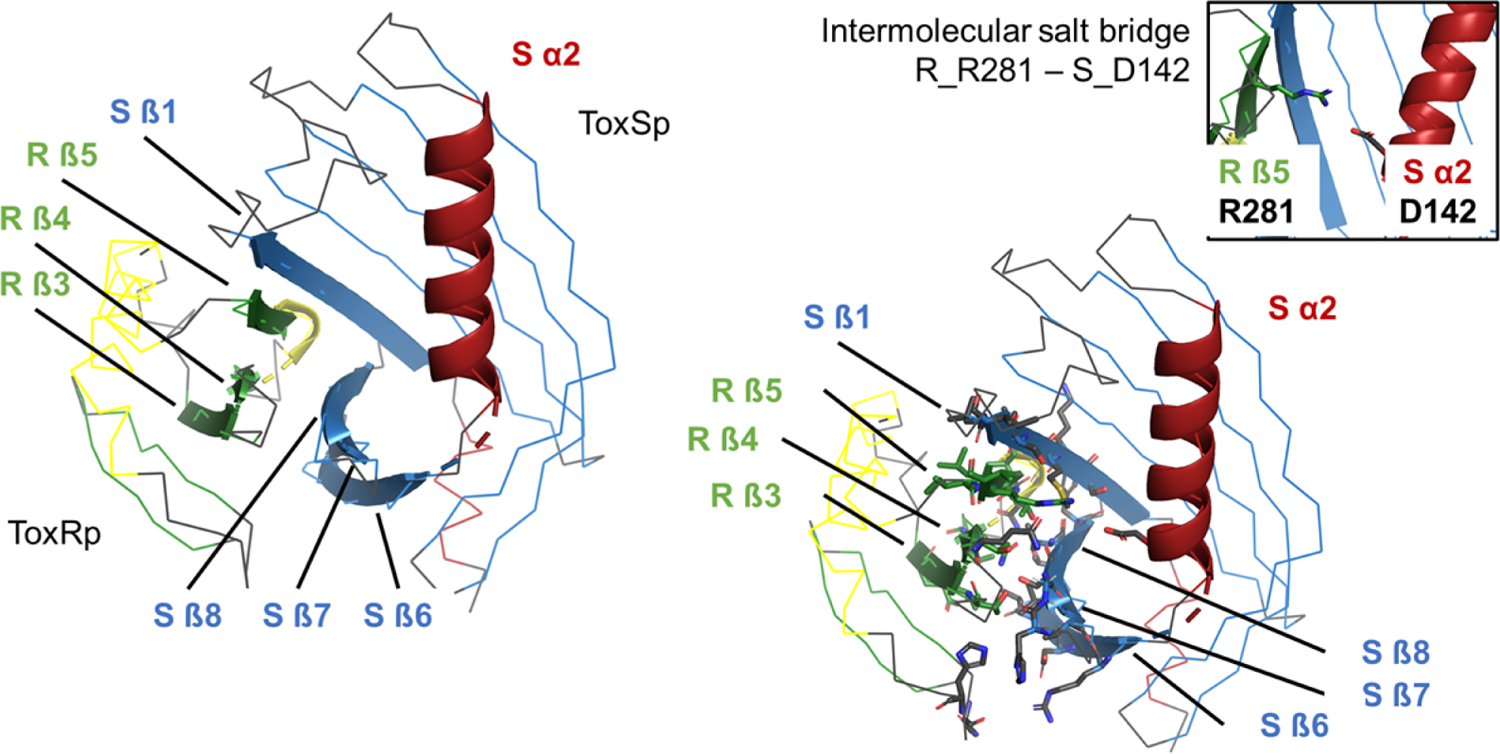
Overview of ToxRSp interface. The interaction is mainly established via ToxRp ß strands 3, 4 and 5, and ToxSp ß strands 6, 7 and 8. Additionally, an intermolecular salt bridge is formed between ToxRp R 281 and ToxSp D142 located at one opening of ToxSp barrel.

**Fig. S3.**
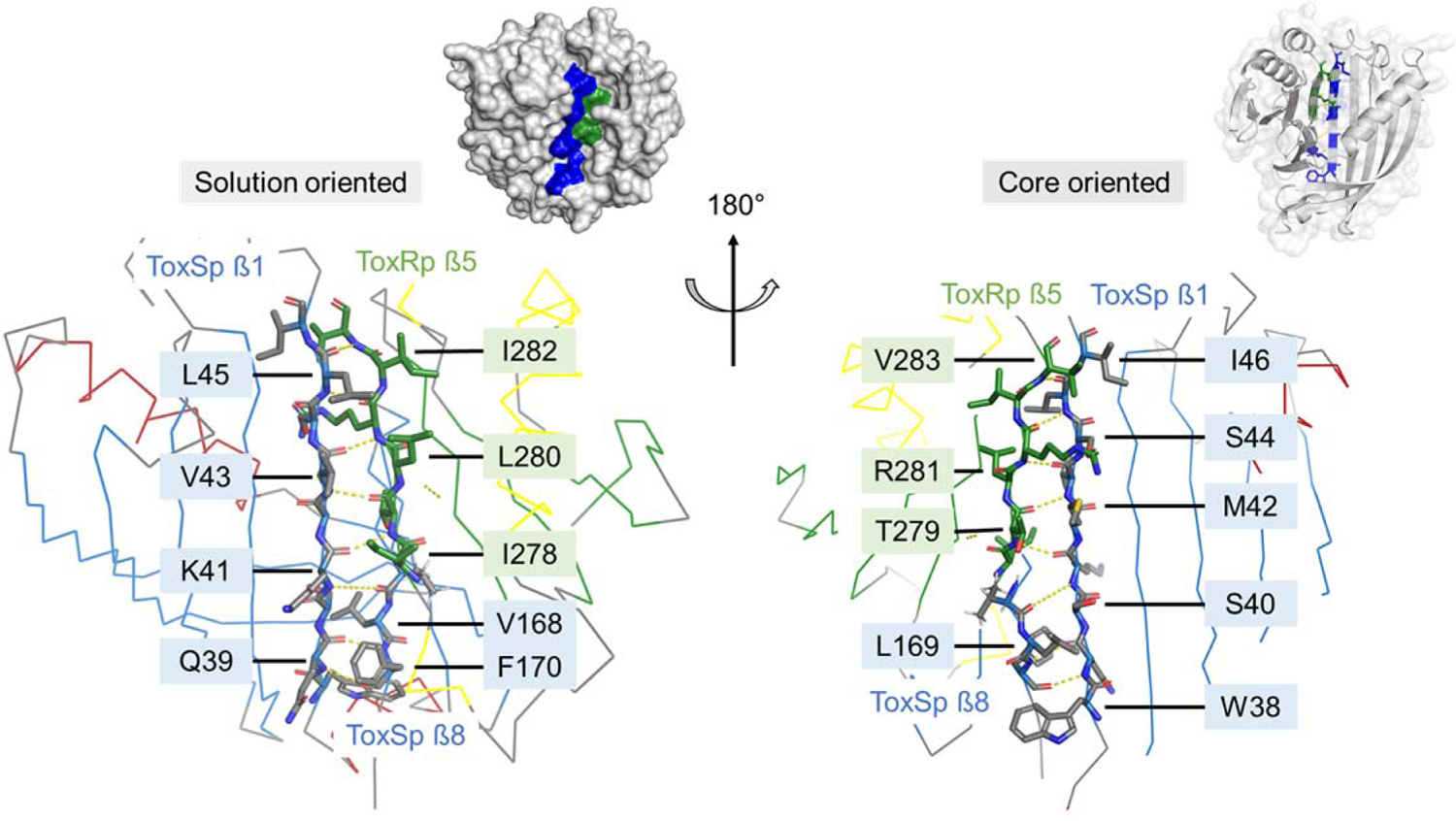
Detailed description of the intermolecular main chain H-bond pattern. H-bond network is established between ToxRp ß5 and ToxSp ß1 thereby filling the gap between ToxSp ß1 and ß8.

**Fig. S4.**
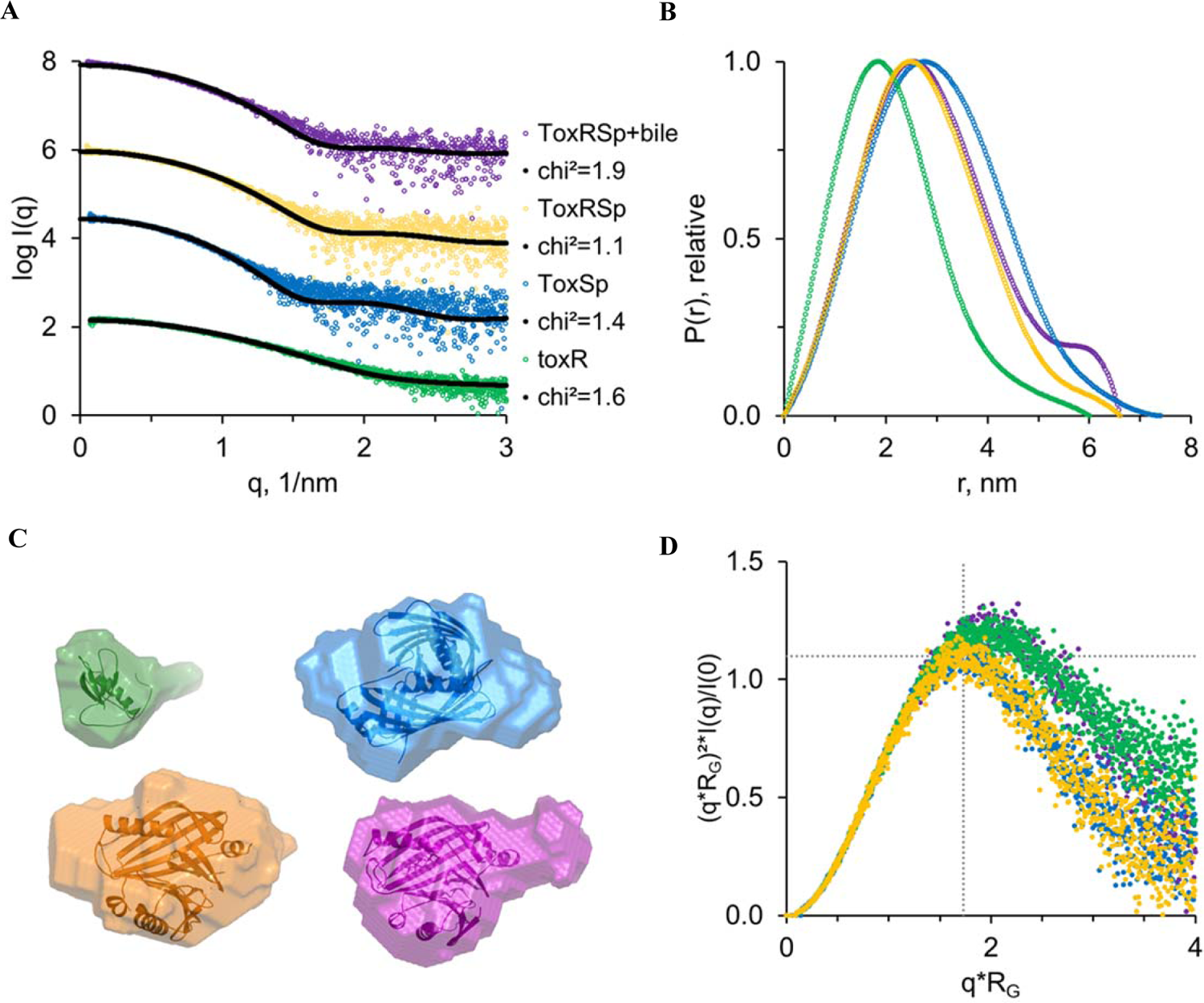
Analysis of size-exclusion chromatography coupled with solution small-angle X-ray scattering SEC-SAXS experiments. ToxRp (green), ToxSp (blue), ToxRSp (orange) and ToxRSp:bile (purple). **A:** The SEC-SAXS profiles are compared to the theoretical scattering curves (Crysol). Chi² values are indicated. Profiles are shifted along the y-axis for clarity. The following models were used ToxRp – NMR structure pdb: 7NN6; ToxSp – the dimeric model (AlphaFold-Multimer); ToxRSp – crystal structure pdb (8ALO); ToxRSp plus bile acid (1:1)-MD complex (Fig. 2). **B:** p(r) functions. **C:** Overlay of the models listed in (A) with the respective bead models calculated from the SAXS profiles. Note, for ToxRp and ToxRSp:bile an excess of volume indicates overall larger size. **D:** Dimensionless Kratkyplot ((qRG)2I(q)/I(0) vs qRG). Note, dotted lines are drawn at y=1.104 and x = √ 3. Folded proteins have a local maximum where the two lines intersect. Here, the maxima of ToxRp and ToxRSp:bile are shifted, indicating a flexible/elongate structure.

**Fig. S5.**
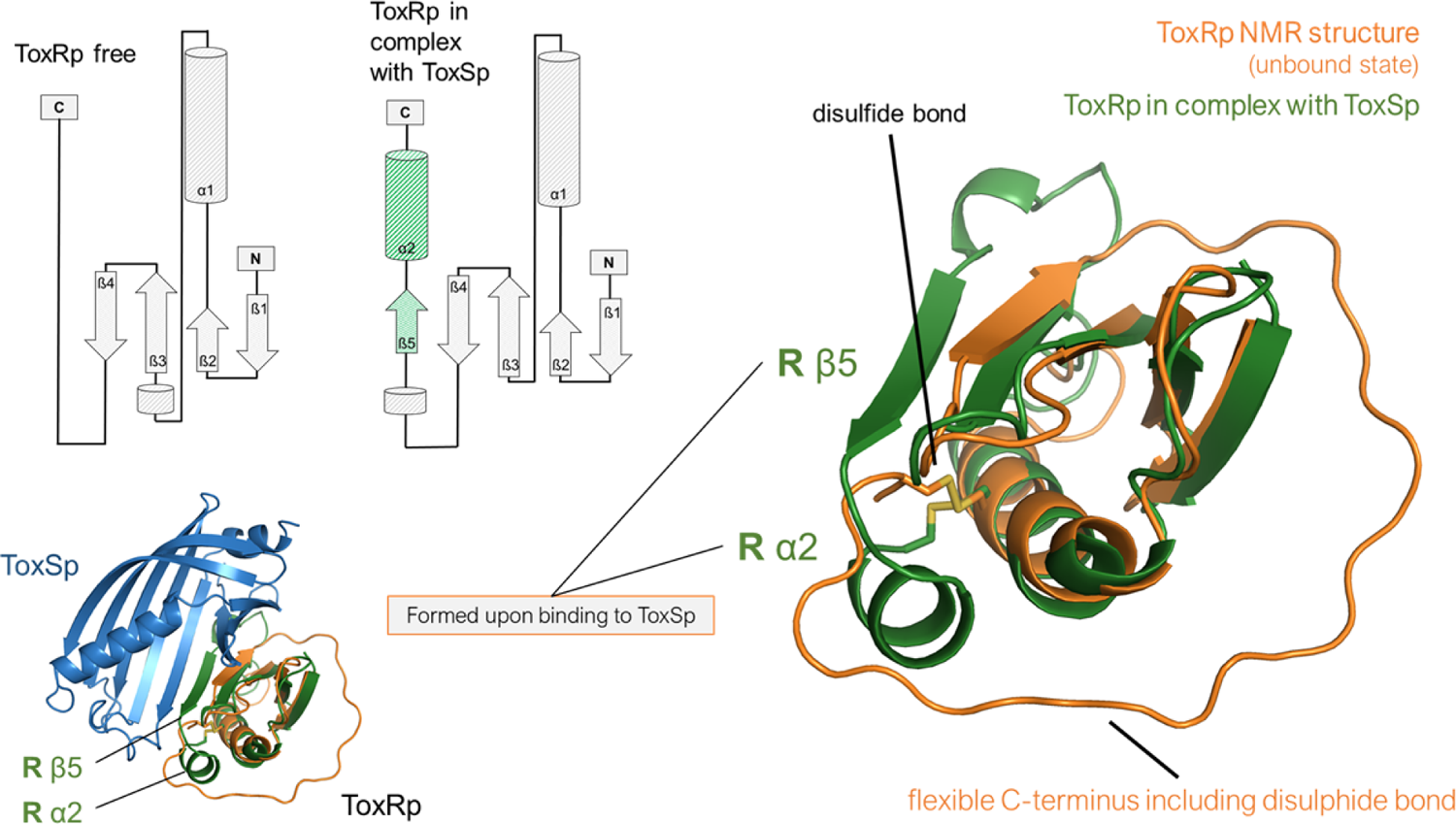
Comparison of free ToxRp and ToxRp bound to ToxSp. Upon binding ToxRp unstructured C-terminal region forms ß5 and α2 which are involved in the establishment of the ToxRSp complex.

**Fig. S6.**
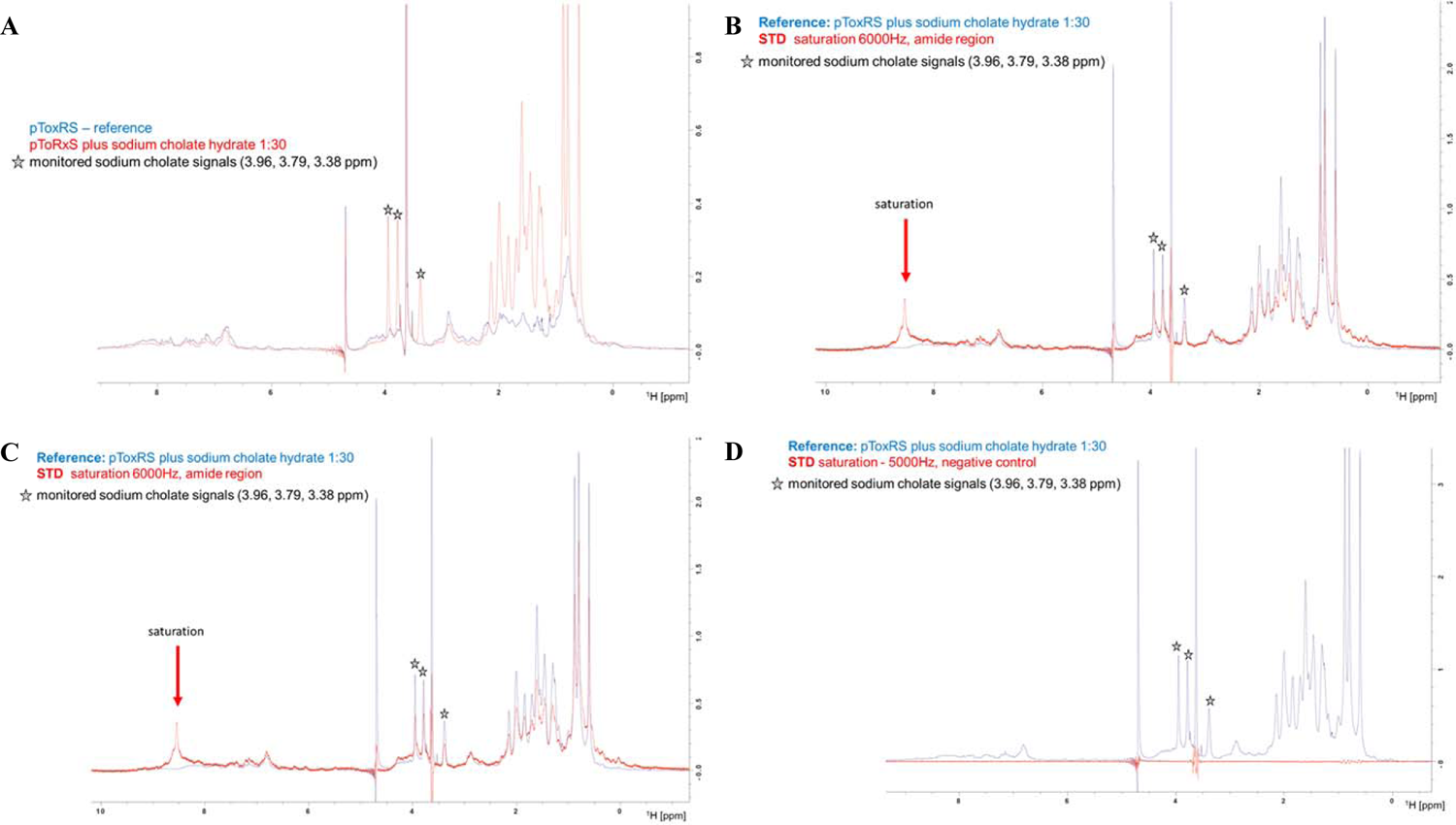
Saturation transfer difference NMR spectra of ToxRSp with sodium cholate hydrate. Spectra clearly reveal a binding between ToxRSp and bile, since saturation in the amide region of ToxRSp resulted in signals of the ligand in the STD spectra.

**Fig. S7.**
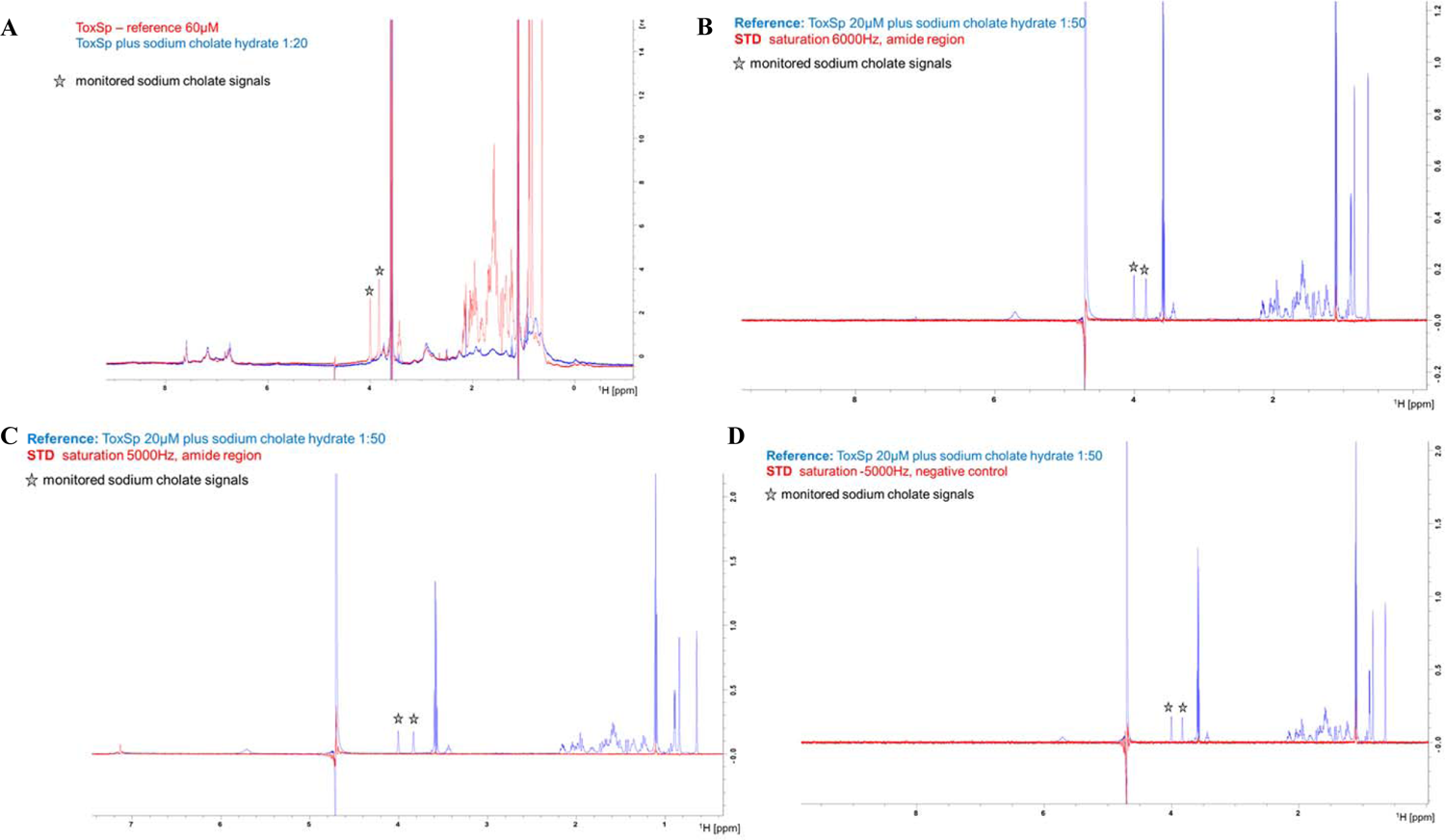
Saturation transfer difference NMR spectra of ToxSp with sodium cholate hydrate. Spectra does not show an interaction between ToxSp and bile. No ligand signals could be observed upon saturation of amide signals of ToxSp.

**Fig. S8.**
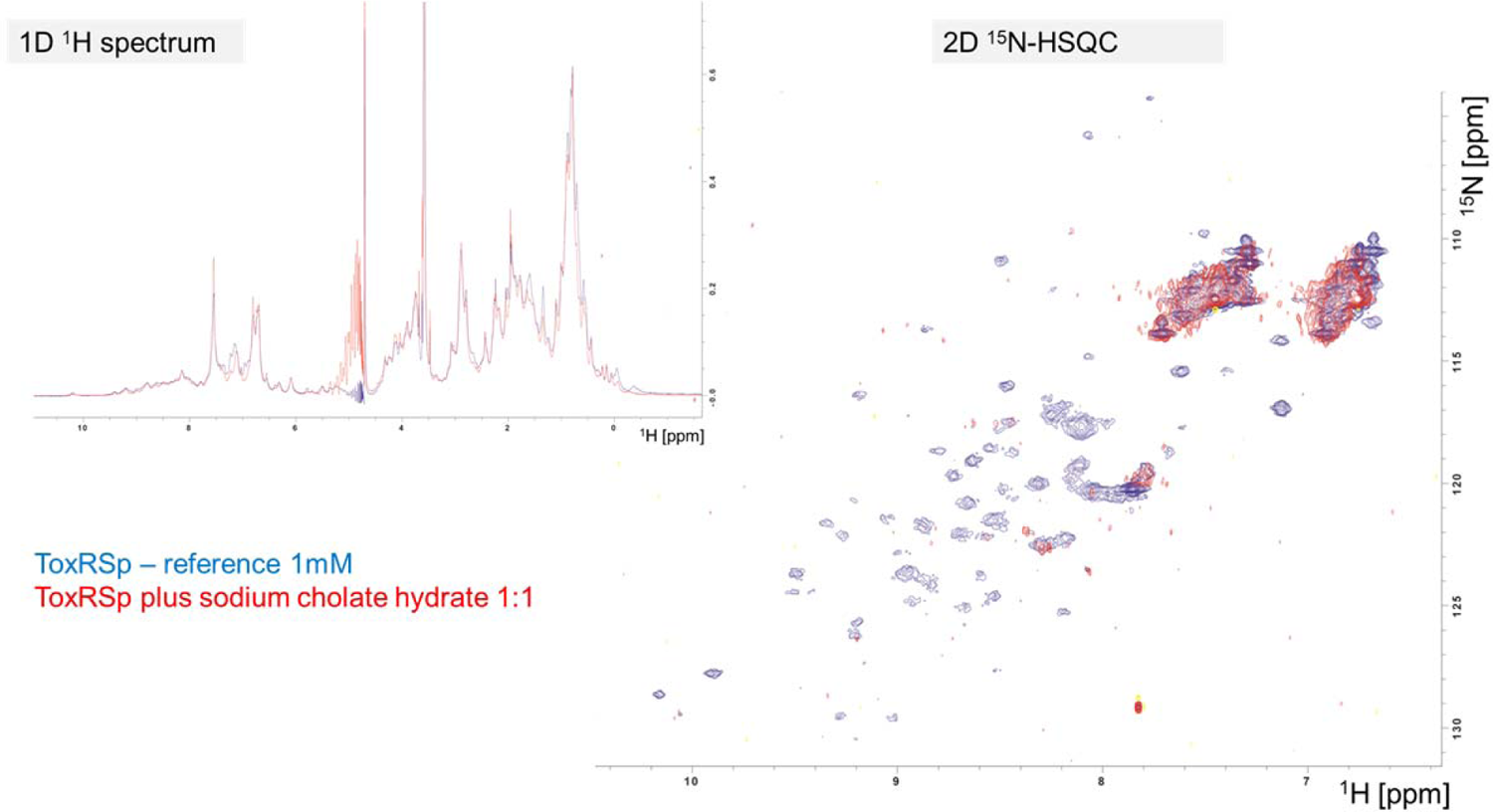
NMR chemical shift perturbance experiments. Comparison of 1D ^1^H spectra of ToxRSp with and without sodium cholate hydrate, clearly shows shifting of the signals of ToxRSp. 2D ^15^N-HSQC experiments propose a binding event with increased molecular weight, since signals of ToxRSp disappear upon bile addition.

**Fig. S9.**
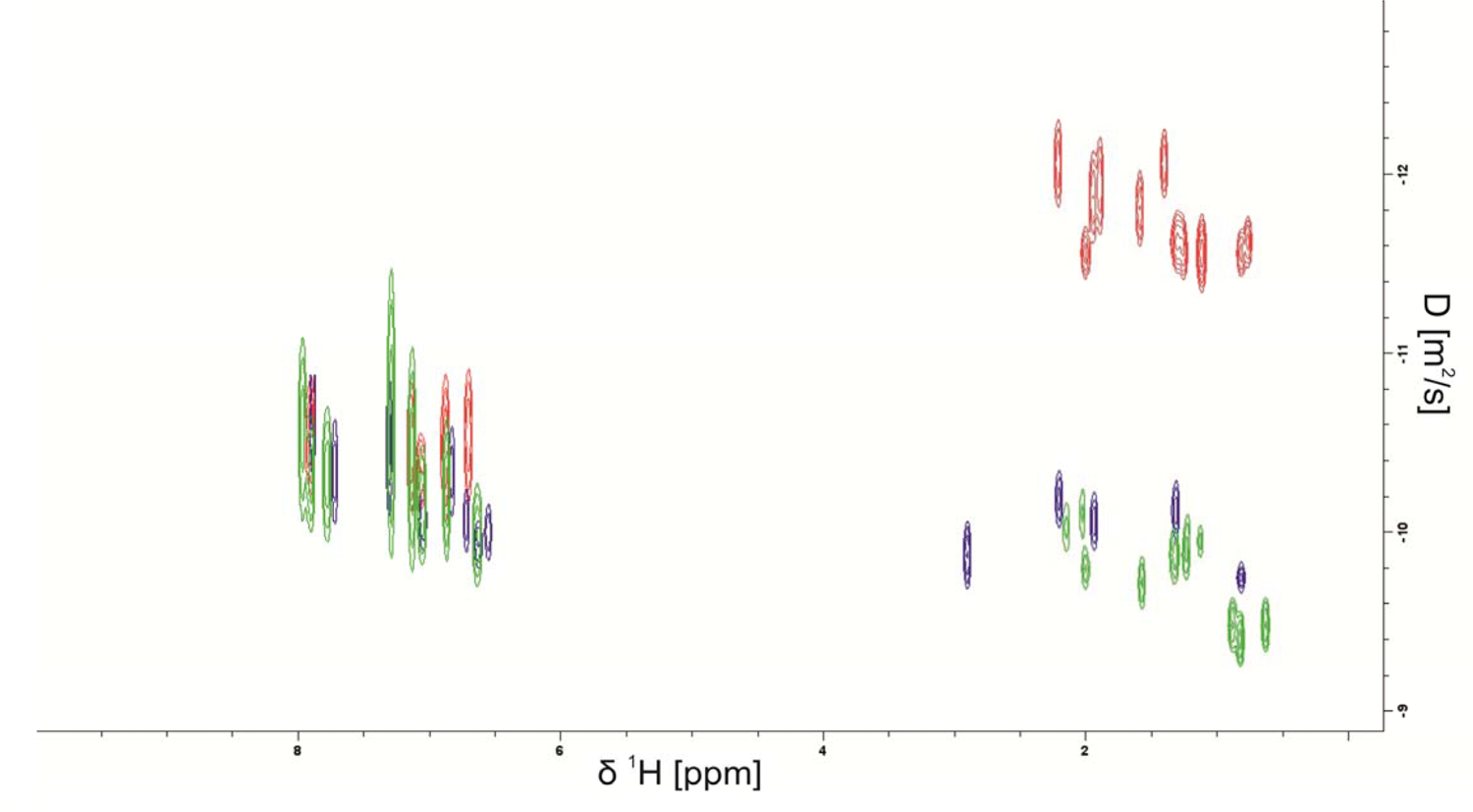
Superimposed NMR DOSY spectra of ToxRSp with and without bile acid sodium cholate hydrate. X-axis represents ^1^H chemical shift, y-axis the determined diffusion coefficient D [m²/s]. Three samples were measured: ToxRSp (blue), ToxRSp 1:1 bile acid (red), ToxRSp 1:20 bile acid (green). Addition of bile acid causes a decrease of the diffusion coefficient due to an increase of molecular weight. Higher bile acid concentrations result in higher diffusion coefficients via averaging due to exchange of bound and free bile molecules.

**Fig. S10.**
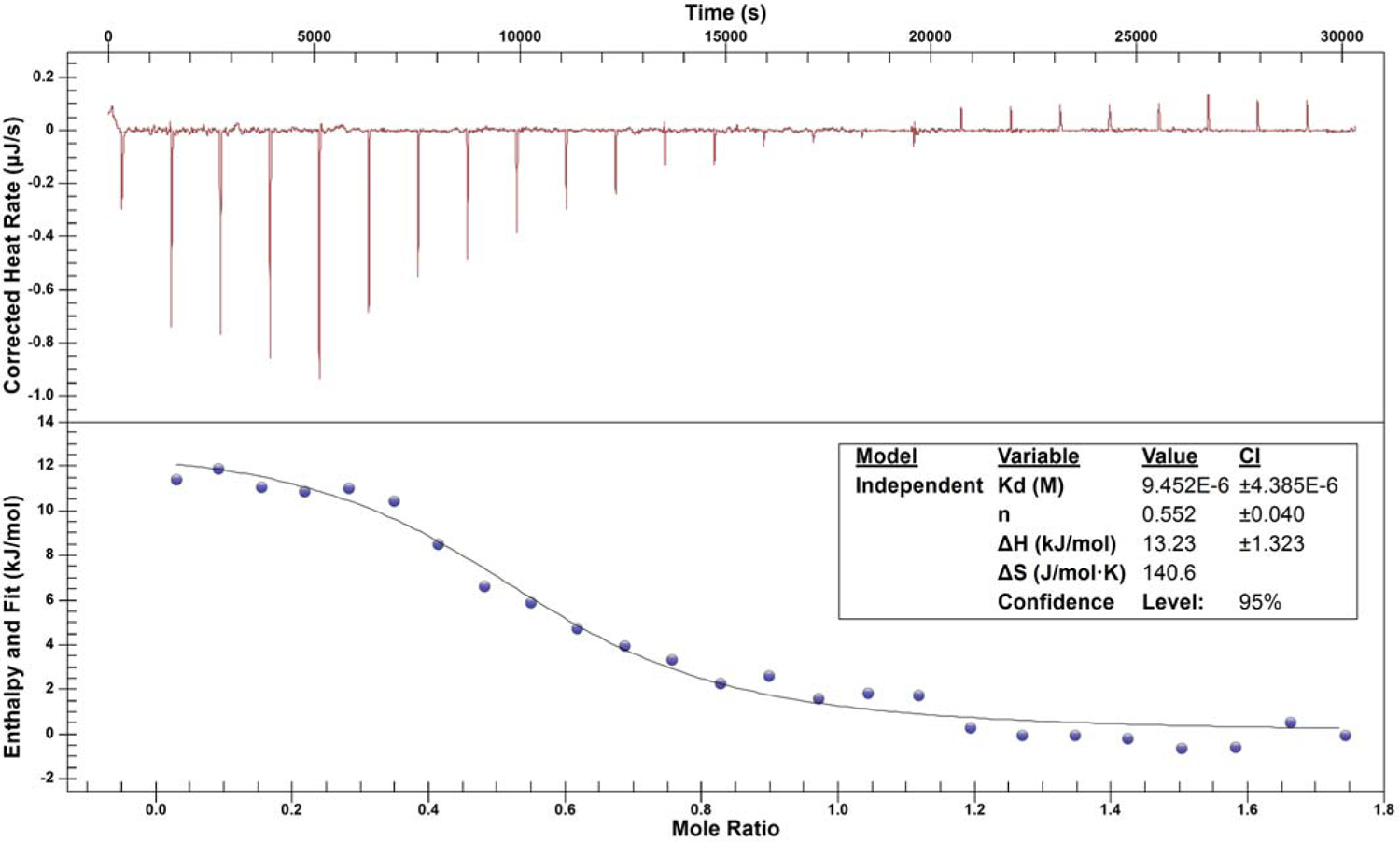
Isothermal titration calorimetry ITC of ToxRSp with sodium cholate hydrate. ITC experiments expose a dissociation constant of 9.5 ± 4.4 µM of ToxRSp and bile acid sodium cholate hydrate.

**Fig. S11.**
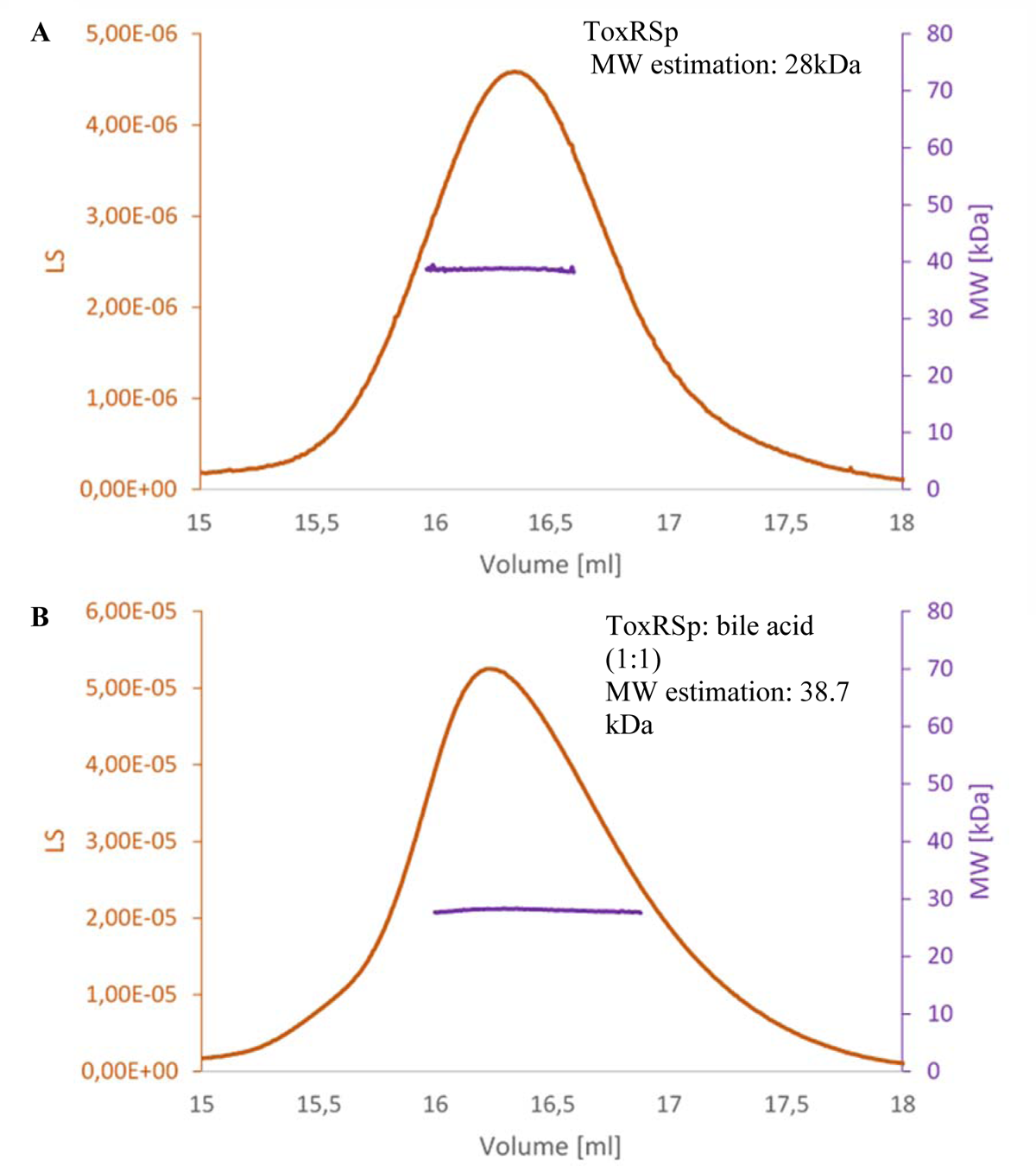
Analysis of size exclusion chromatography with multi-angle static light scattering SEC-MALS. The light scattering curve of the SEC MALS run showing the elution peaks of ToxRS. Light scattering LS is shown in orange and molecular weight MW is depicted in purple. **A:** elution peak of ToxR mixed with bile salt 1:1. **B:** elution peak of ToxRS without bile acid sodium cholate hydrate. Both samples elute in a single peak with a retention time of 16.4 ml for ToxRS with bile acid and 16.3 ml for ToxRS without bile acid. The hydrodynamic radius of ToxRS with bile salts determined by the MALS detector leads to a molecular weight of 38.7 kDa and a hydrodynamic radius of 10.8 nm. The determined molecular weight by SEC-MALS of ToxRSp alone is 28 kDa. These results show an increase of molecular weight upon bile addition.

**Fig. S12.**
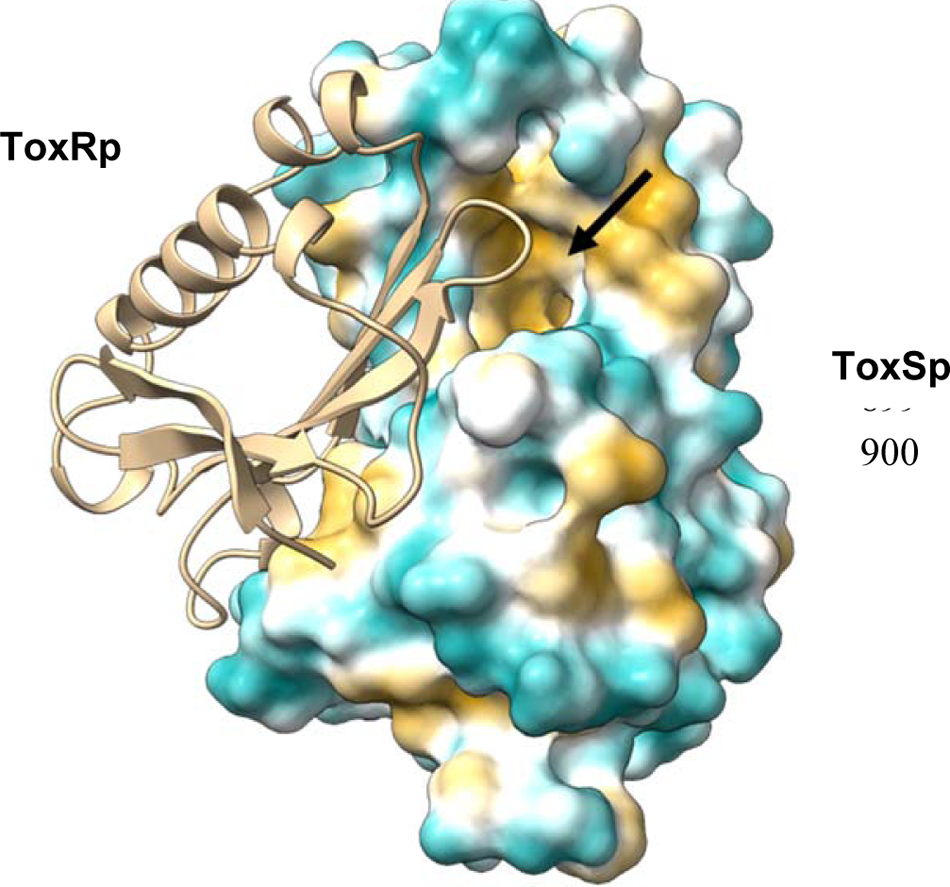
Hydrophobic cleft of ToxSp binding pocket in complex with ToxRp. Hydrophobic (ochre) and hydrophilic (teal) surface representation of ToxS calcualted with ChimeraX mlp. The bile binding cavity is highly hydrophobic (arrow).

**Fig. S13.**
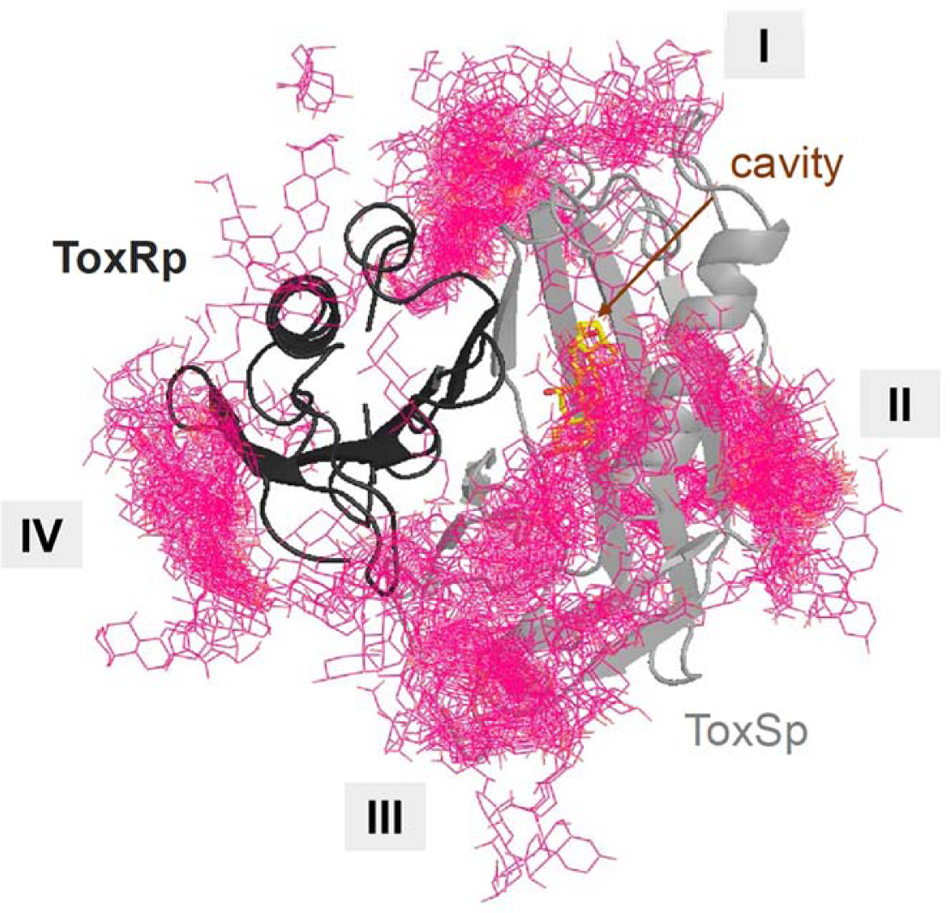
Surface exposed bile interacting regions of ToxRSp determined via MD. Representative structure of the ToxRSp complex (dark grey/light grey cartoons) with cholate bound at the cavity (sticks, C-atoms in yellow) and superimposition of the closest cholate molecules (shown as red lines) detected via MD simulations. Besides the bile binding cavity of ToxSp, four regions (marked as I-IV) on the ToxRSp surface could be mapped as additional bile binding areas. Region I and III involves both proteins, whereas region II is located on ToxSp and region IV on ToxRp.

**Fig. S14.**
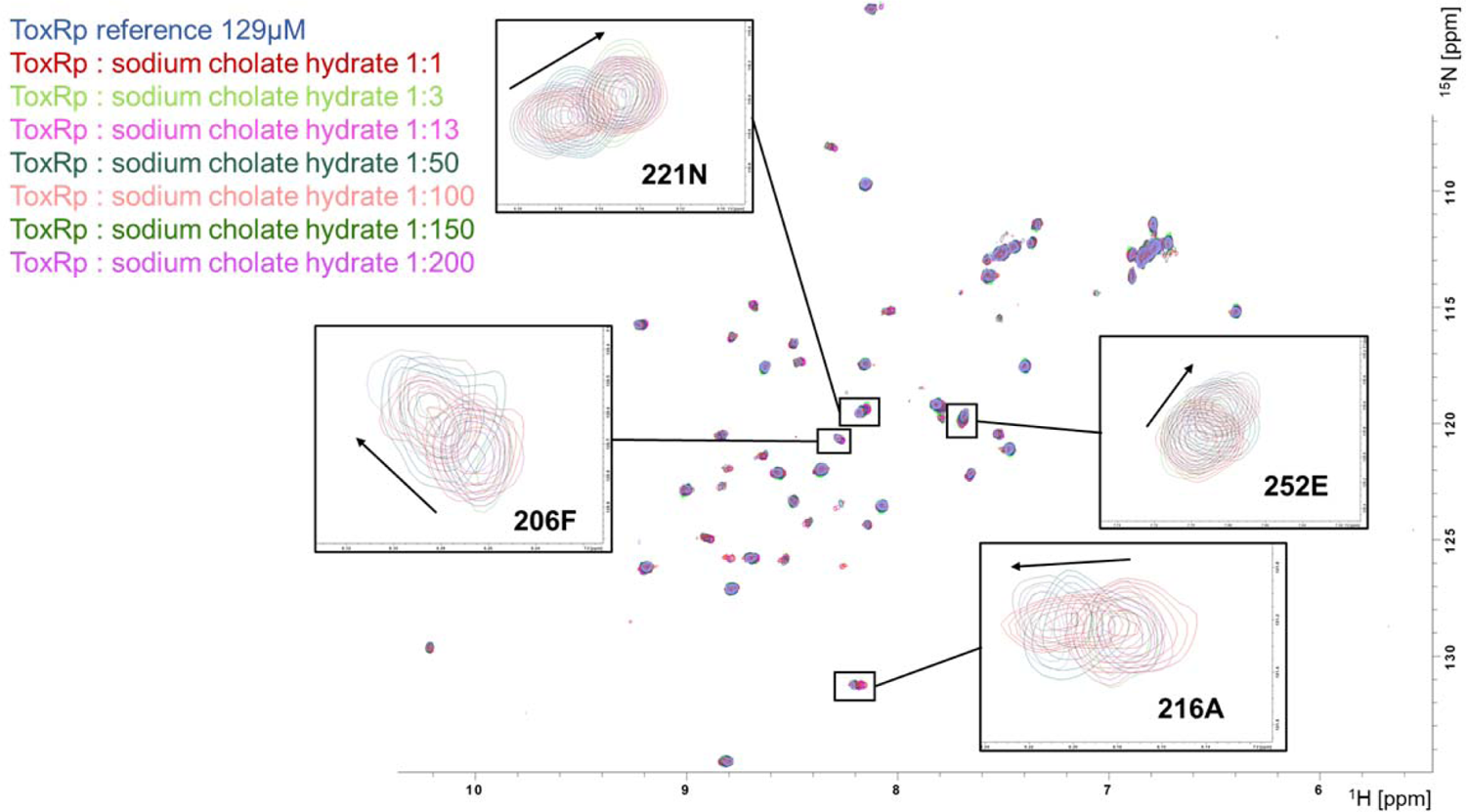
NMR titration experiment with ToxRp and sodium cholate hydrate. Overlay of ^15^N-HSQC spectra of ToxRp with increasing amounts of bile propose a weak interaction event. Significant chemical shift changes of ToxRp could be observed with high ligand concentrations of 1:50-1:200.

**Fig. S15.**
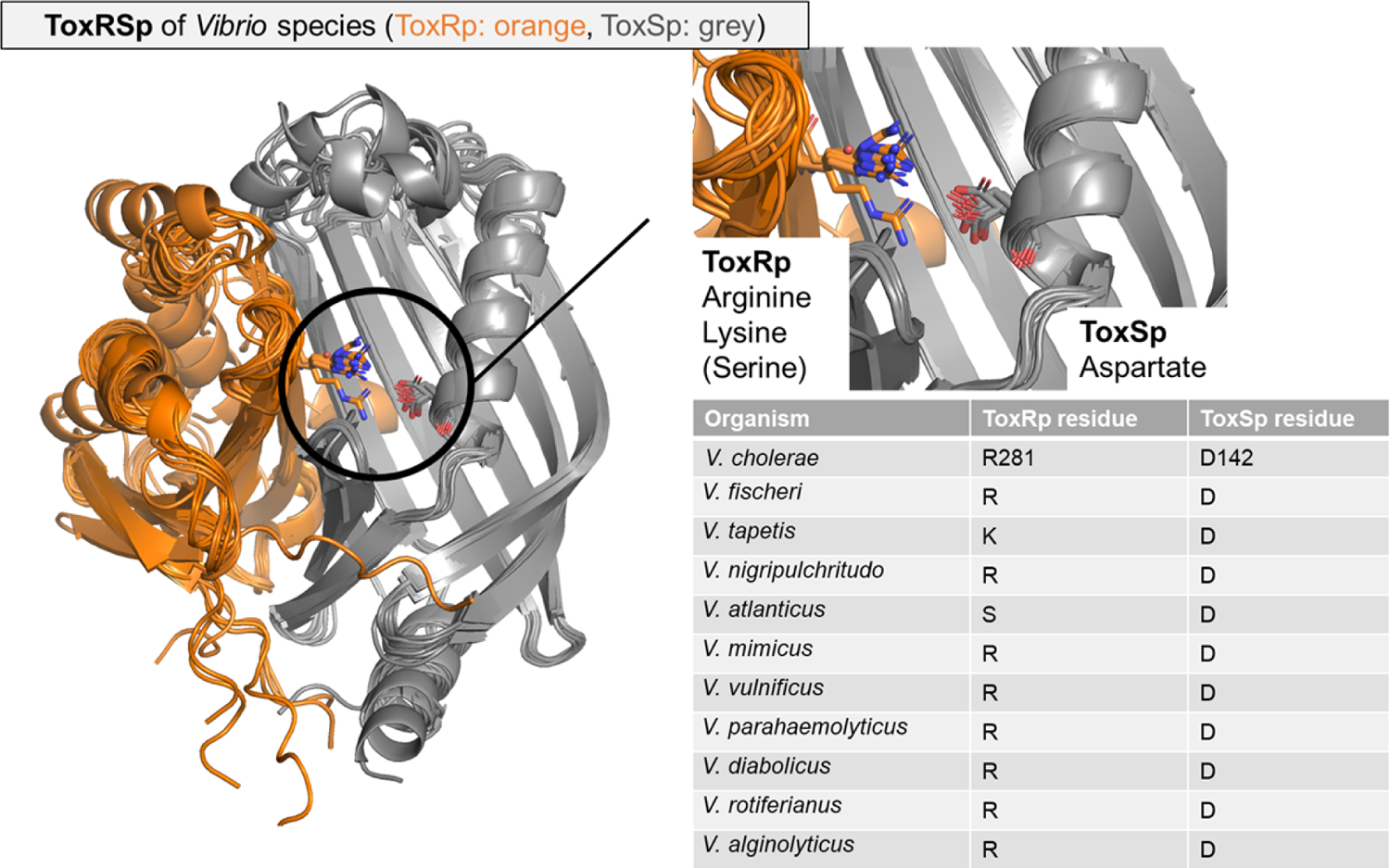
Overlay of ToxRSp from different *Vibrio* species. Structures are calculated using AlphaFold-Multimer (*Evans et al., 2021; Jumper et al., 2021*). The structural alignment reveals a conserved fold of ToxSp (grey), whereas ToxRp (orange) varies among different *Vibrio* strains. The intermolecular salt bridge between ToxRp and ToxSp is highlighted, and residues types involved are mentioned in the table beside the aligned structures. ToxSp salt bridge forming aspartate residue is strictly conserved.

**Fig. S16.**
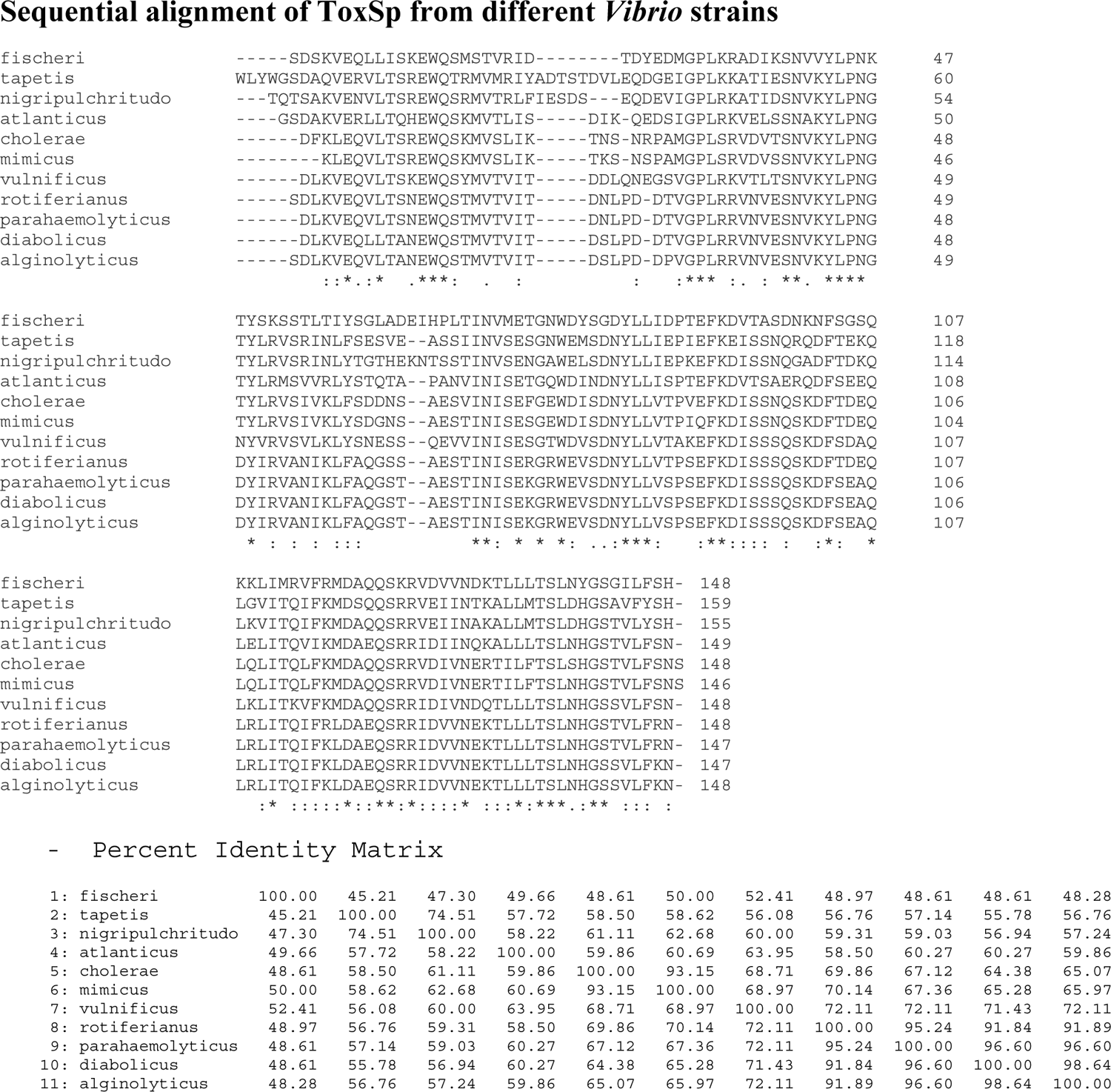
Sequential analysis of ToxSp proteins from different *Vibrio* species. Analysis was performed with Clustal2.1 (*Madeira et al., 2022*). Strongly conserved residues are marked with an asterisk (*), conserved residues with colon (:) and period (.). Below the sequence identities are shown in percent.

**Fig. S17.**
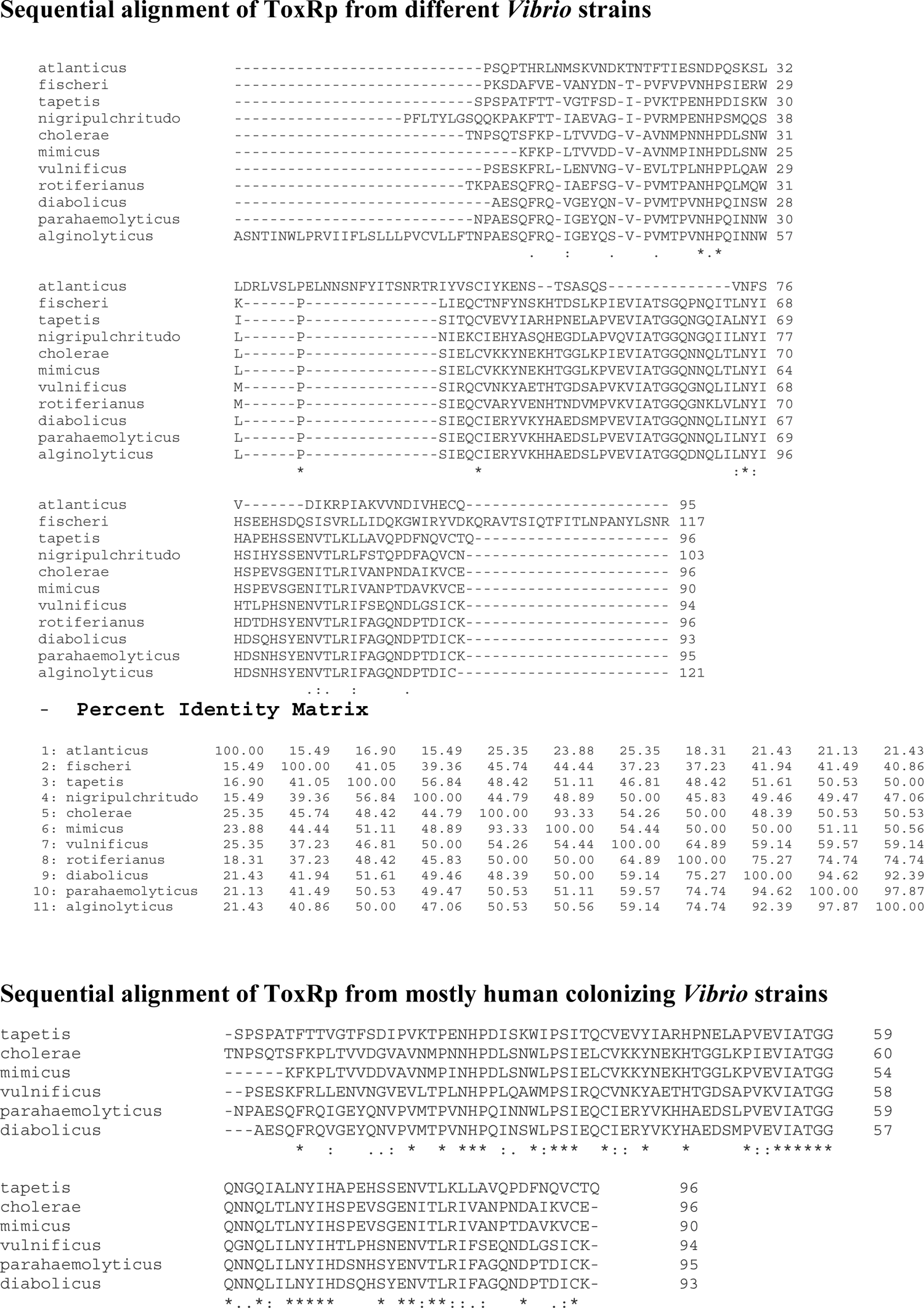
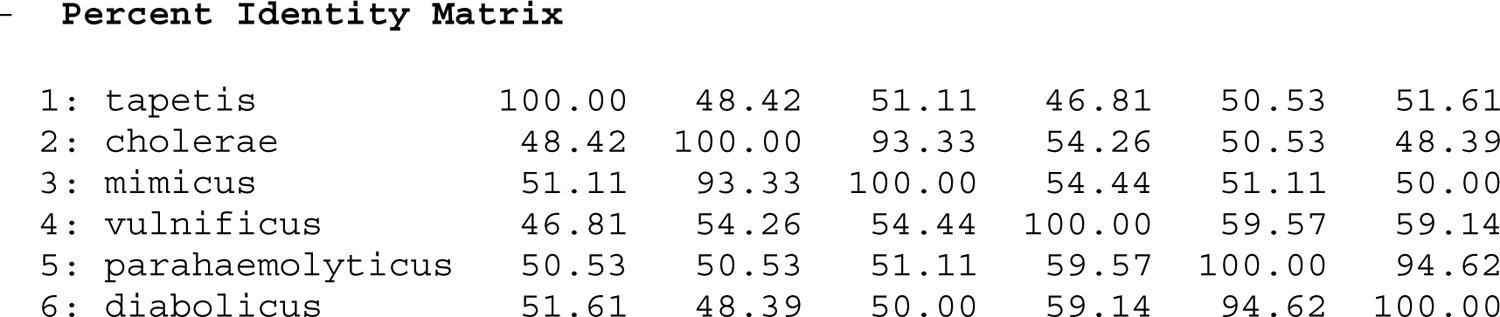
Sequential analysis of ToxRp proteins from different *Vibrio* species. Analysis was performed with Clustal2.1 (*Madeira et al., 2022*). Strongly conserved residues are marked with an asterisk (*), conserved residues with colon (:) and period (.). Below the sequence identities are shown in percent. Due to the higher variability of ToxRp in *Vibrio’s* an additional analysis was performed for closely related strains, which are mostly human colonizing.

**Fig. S18.**
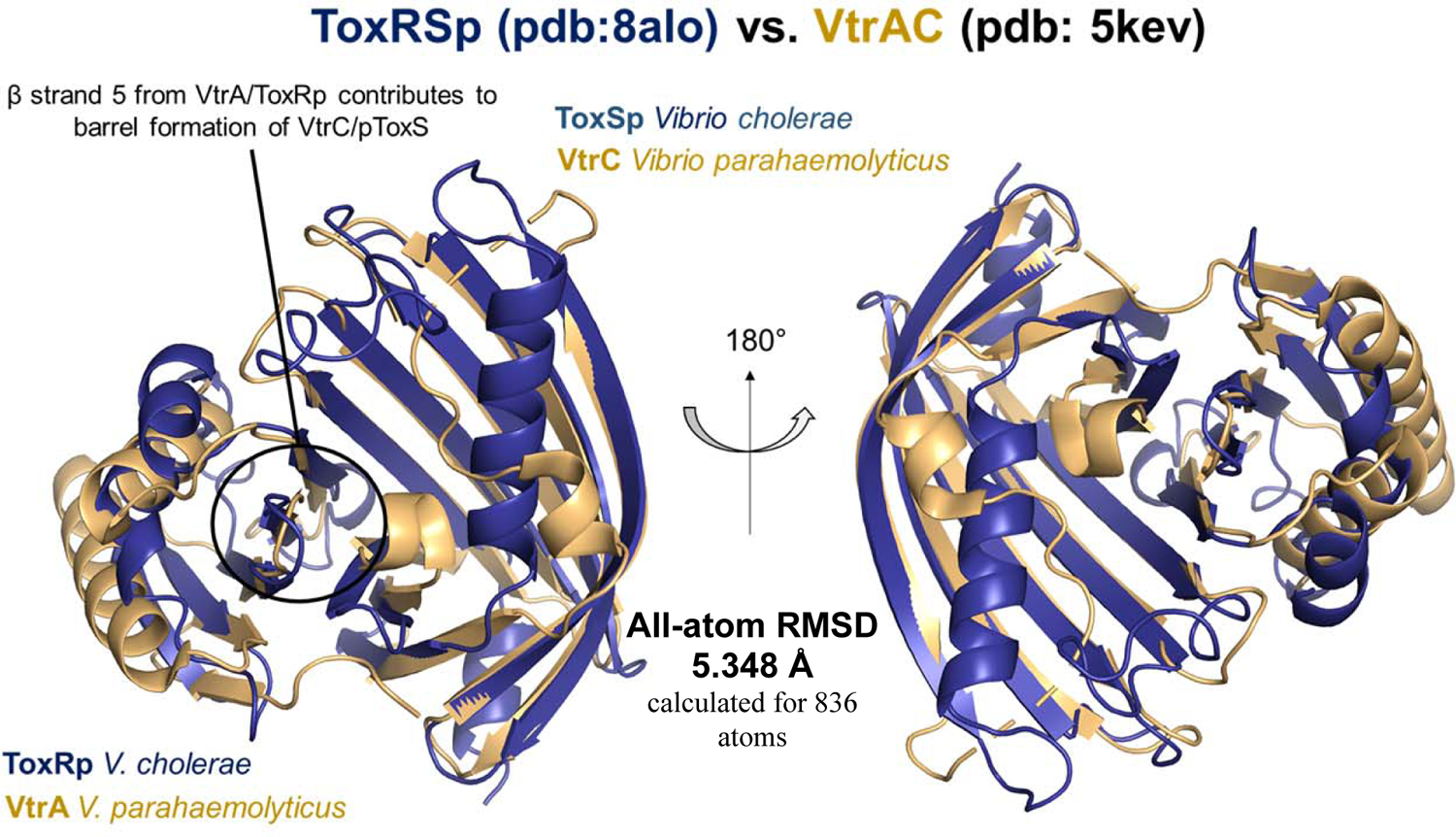
Structural alignment of bile interacting protein complexes ToxRS *V. cholerae* and VtrAC *V. parahaemolyticus*. VtrAC (gold; pdb: 5KEV) and ToxRSp (dark blue; pdb: 8ALO) reveal a similar fold (RMSD: 3.8Å). VtrC and ToxSp form a ß-barrel fold, which is stabilized by ß-strand 5 from VtrA and ToxRp, resulting in an obligate heterodimer formation including a binding pocket in the barrel cavity.

**Table S1.**
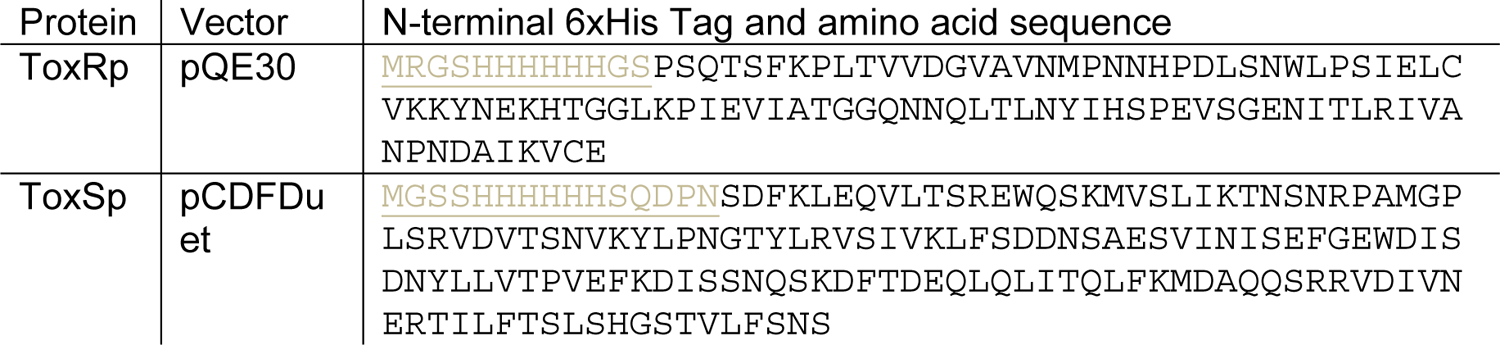
Description of expression constructs. List of plasmids and amino acid sequences used for the expression of ToxRp and ToxSp.

**Table S2.** Crystal data and structure refinement table of ToxRSp.

**ToxRSp crystal structure**
|  |  |
| --- | --- |
| Resolution range | 49.27 - 3.002 (3.109 - 3.002) |
| Space group | P 6<sub>5</sub> |
| Unit cell (Å, °) | 72.507, 72.507, 79.463<br>90, 90, 120 |
| Total reflections | 9392 (909) |
| Unique reflections | 4786 (467) |
| Multiplicity | 2.0 (1.9) |
| Completeness (%) | 99.85 (100.00) |
| Mean I/sigma(I) | 13.10 (2.18) |
| Wilson B-factor | 75.07 |
| R-merge | 0.03888 (0.3675) |
| R-meas | 0.05498 (0.5197) |
| R-pim | 0.03888 (0.3675) |
| CC1/2 | 0.999 (0.591) |
| CC* | 1 (0.862) |
| Reflections used in refinement | 4785 (467) |
| Reflections used for R-free | 260 (38) |
| R-work | 0.2163 (0.2886) |
| R-free | 0.2491 (0.3463) |
| CC(work) | 0.940 (0.656) |
| CC(free) | 0.884 (0.774) |
| Number of non-hydrogen atoms | 1865 |
| Macromolecules | 1854 |
| Ligands | 0 |
| Solvent | 11 |
| Protein residues | 236 |
| RMS(bonds) | 0.012 |
| RMS(angles) | 1.86 |
| Ramachandran favored (%) | 93.53 |
| Ramachandran allowed (%) | 6.47 |
| Ramachandran outliers (%) | 0.00 |
| Rotamer outliers (%) | 0.92 |
| Clashscore | 4.32 |
| Average B-factor | 87.41 |
| Macromolecules | 87.60 |
| Solvent | 55.04 |

**Table S3.**
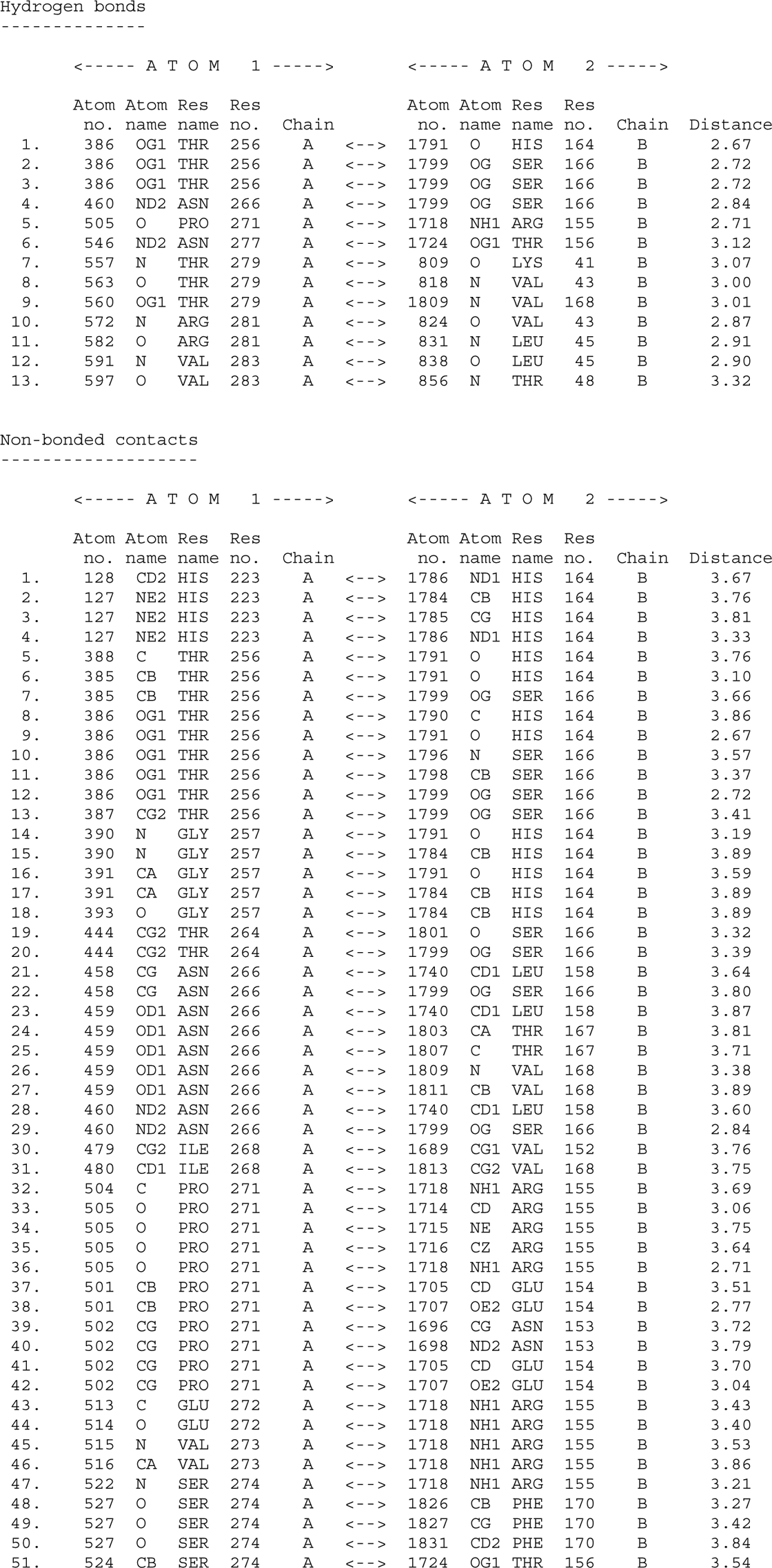
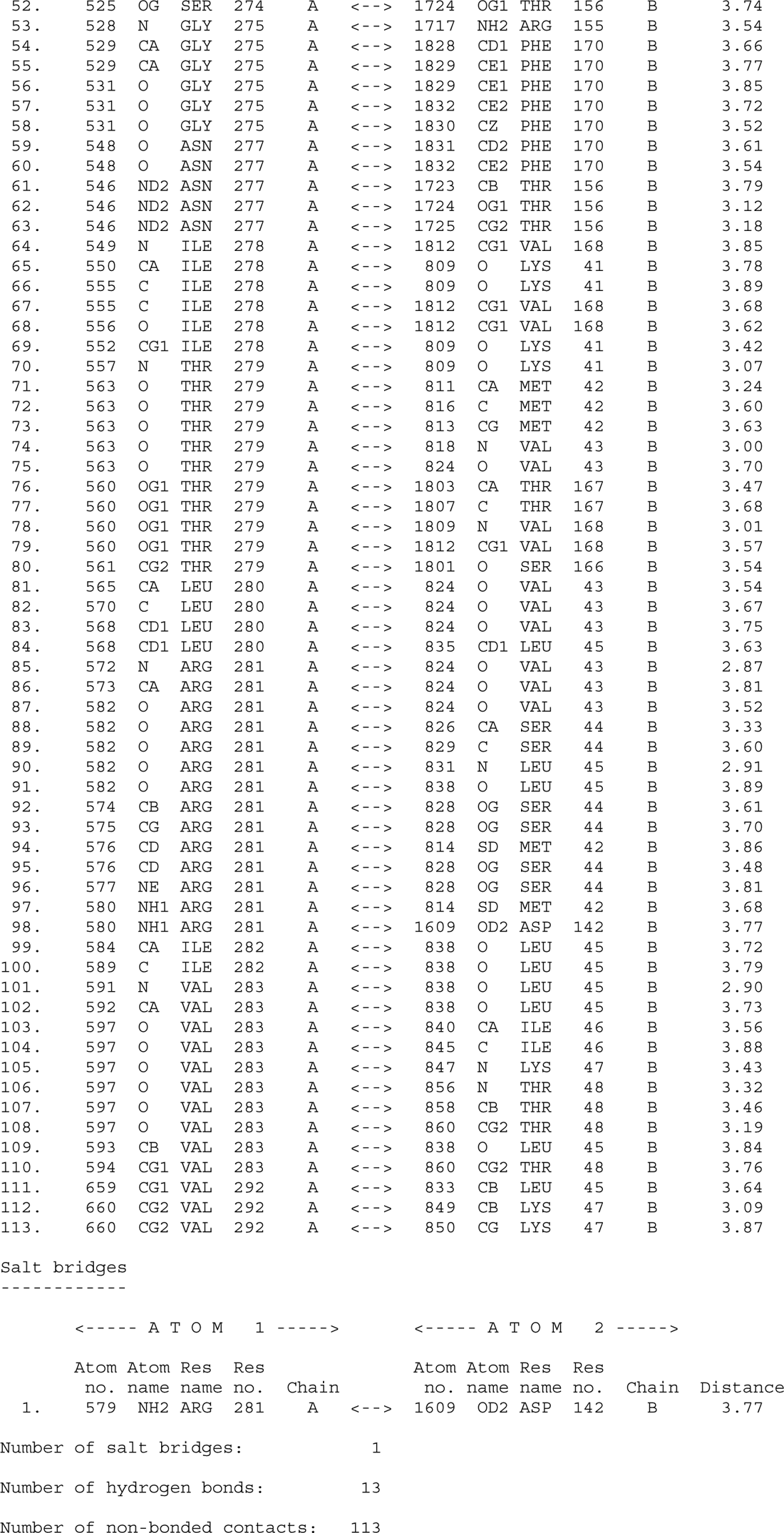
List of all atom interactions of ToxRSp (pdb: 8ALO). The list was created using PDBsum online server (; *Laskowski et al., 2018*).

**Table S4.** Analysis of size-exclusion chromatography coupled with solution small-angle X-ray scattering SEC-SAXS. According to SAXS curves, ToxRp is monomeric in solution, whereas ToxSp seems to form dimers. Upon bile addition the molecular weight of the ToxRSp complex increases.

|  | Radius of Gyration,<br>R<sub>G</sub> [nm] | Dmax<br>[nm] | Chi<sup>2</sup> | MW estimation [kDa] |  |  |
| --- | --- | --- | --- | --- | --- | --- |
|  |  |  |  | MoW | VC | Bayesian |
| ToxRp | 1.6 +/- 0.1 | 6.0 +/- 0.2 | 1.6 | 9 +/- 1 | 11 +/- 1 | 10 +/- 1 |
| ToxSp<br>(dimeric) | 2.3 +/- 0.1 | 7.4 +/- 0.2 | 1.4 | 33 +/- 3 | 35 +/- 3 | 35 +/- 3 |
| ToxRSp | 2.1 +/- 0.1 | 6.6 +/- 0.2 | 1.1 | 30 +/- 3 | 28 +/- 3 | 28 +/- 3 |
| ToxRSp + bile<br>(1:1) | 2.4 +/- 0.1 | 6.6 +/- 0.5 | 1.9 | 32 +/- 3 | 29 +/- 3 | 29 +/- 3 |

**Table S5.**
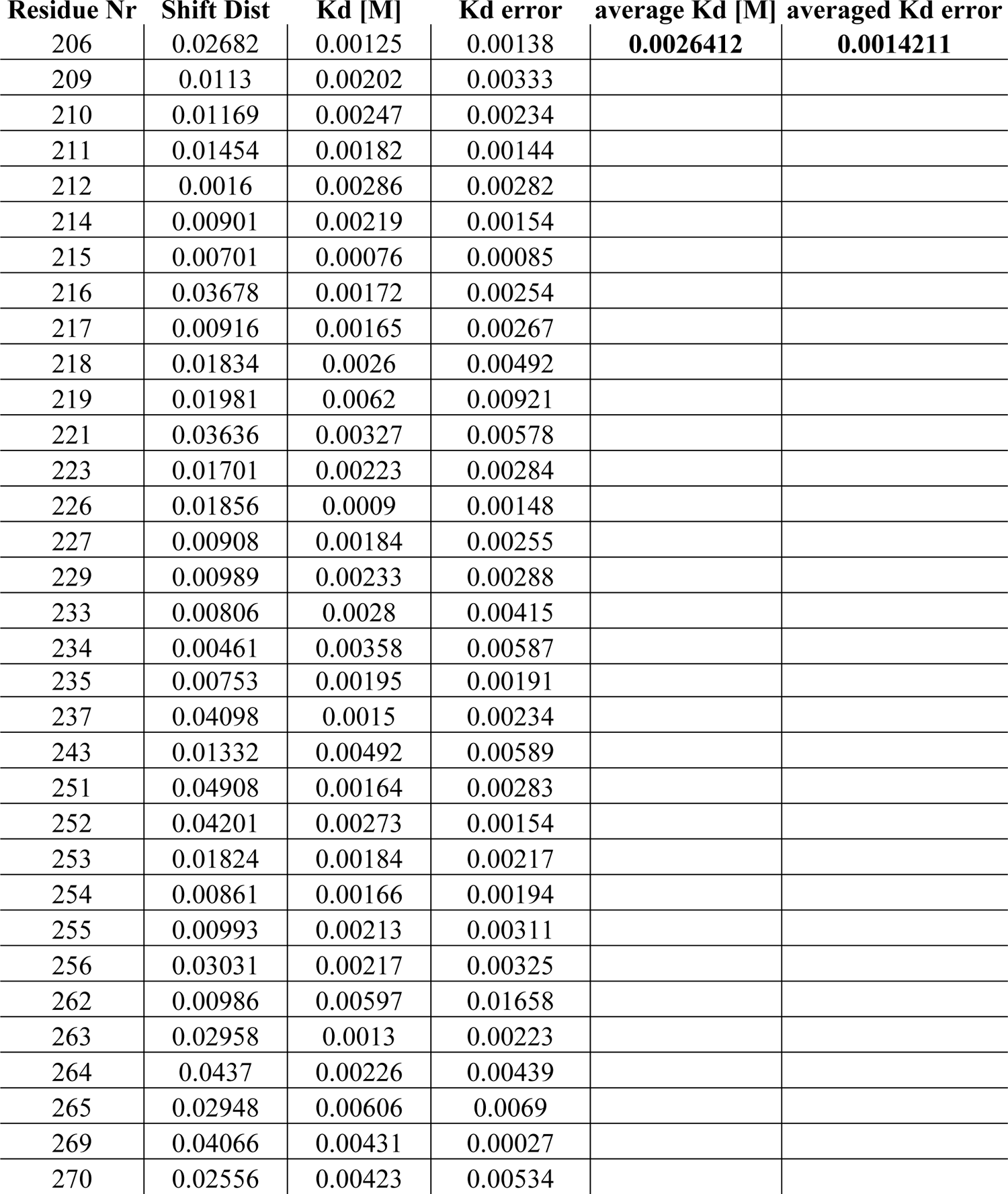
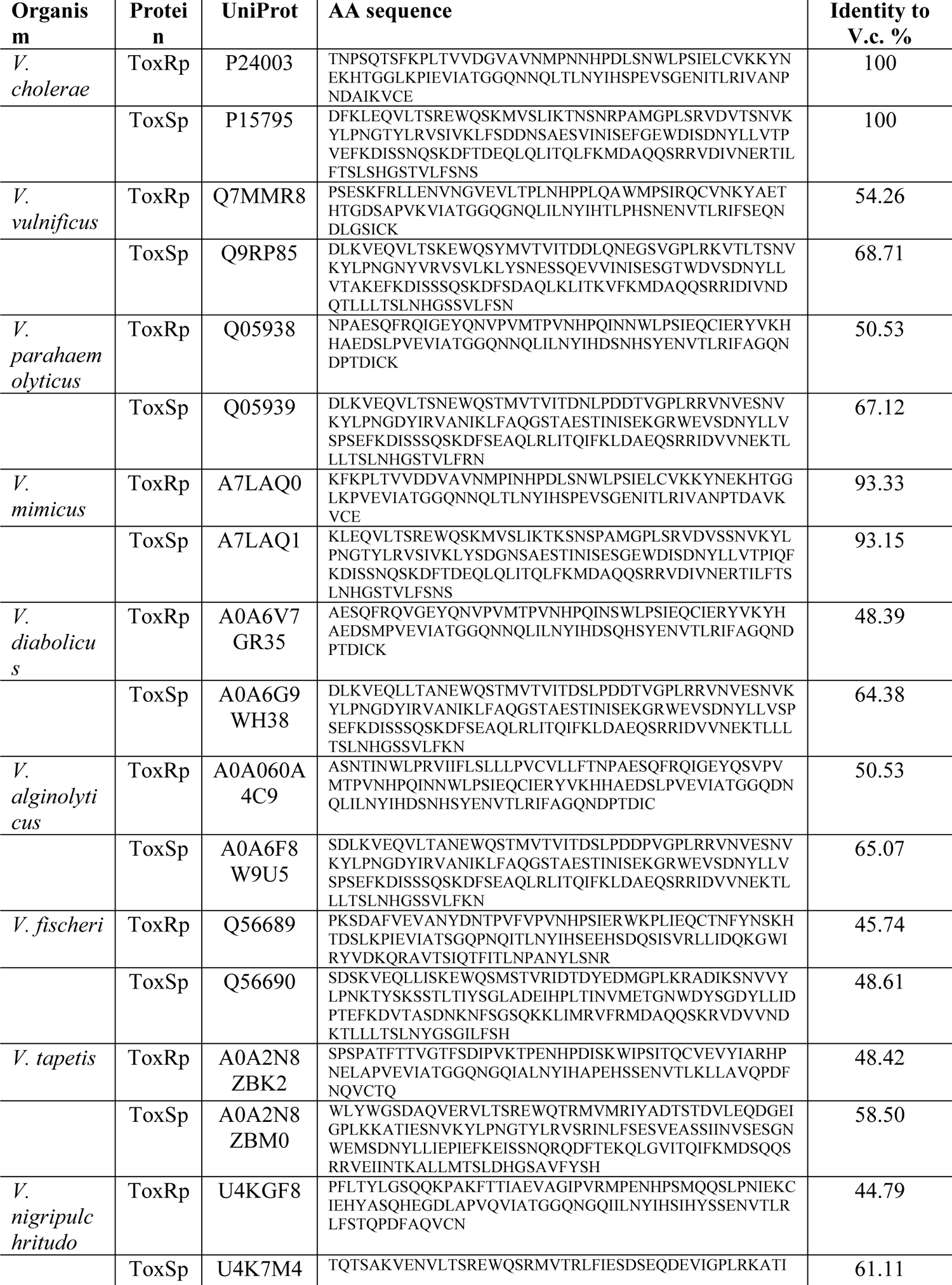
NMR derived dissociation constants for ToxRp and sodium cholate hydrate. NMR titration experiments reveal a weak binding of bile to ToxRp with a dissociation constant of 2.6 mM. Table S5 includes ToxRp residue number, chemical shift distance, dissociation constant Kd [M], error of Kd. Last rows are the calculated average Kd as well as the averaged error.

**Table S6.**
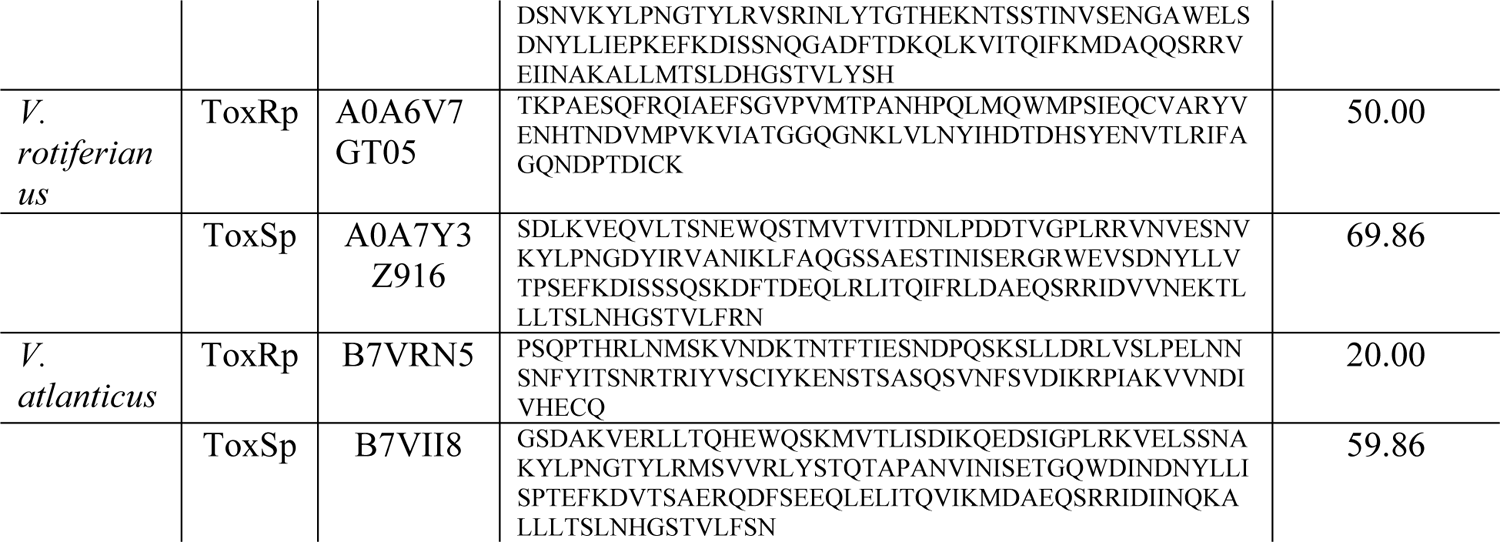
List of amino acid sequences of sensory domains of ToxRS proteins from different Vibrio species. Table S5 includes UniProt ID of the used sequences as well as sequence identity values to ToxRSp from *V. cholerae*.

**Table S7.** Structural homology search of ToxSp bound to ToxRp using Dali server (*Holm, 2022*).

|  |  |  |  |  |  |  |  |
| --- | --- | --- | --- | --- | --- | --- | --- |
| 1 | 5KEW-B | 12 | 2.6 | 112 | 132 | 10 | VTRA PROTEIN |
| 2 | 4zgf-A | 9.6 | 3.6 | 112 | 141 | 13 | UNCHARACTERIZED PROTEIN |
| 3 | 2kts-A | 8.4 | 2.9 | 97 | 117 | 7 | HEAT SHOCK PROTEIN HSLJ |
| 4 | 4l3r-B | 8.4 | 3.2 | 109 | 144 | 5 | UNCHARACTERIZED PROTEIN |
| 5 | 4u3q-B | 7.9 | 1.9 | 84 | 99 | 17 | 17 KDA LIPOPROTEIN |
| 6 | 2mhd-A | 7.6 | 2.7 | 91 | 110 | 13 | UNCHARACTERIZED PROTEIN |
| 7 | 4iab-A | 7.5 | 4.3 | 97 | 142 | 9 | HYPOTHETICAL PROTEIN |
| 8 | 6ohh-B | 7.5 | 2.7 | 98 | 128 | 12 | EF1P2_MFAP2B |
| 9 | 5byp-A | 7.4 | 3.1 | 91 | 121 | 13 | PUTATIVE UNCHARACTERIZED PROTEIN |
| 10 | 3lhn-A | 7.3 | 2.6 | 91 | 108 | 9 | LIPOPROTEIN |
| 11 | 3hty-A | 7.2 | 3.1 | 82 | 94 | 9 | HYPOTHETICAL PROTEIN BT_0869 |
| 12 | 4n7c-A | 7.2 | 2.4 | 95 | 174 | 7 | BLA G 4 ALLERGEN VARIANT 1 |
| 13 | 2la7-A | 7.2 | 2.5 | 99 | 145 | 8 | UNCHARACTERIZED PROTEIN |
| 14 | 2erv-A | 7.1 | 3.3 | 99 | 150 | 6 | HYPOTHETICAL PROTEIN PAER03002360 |
| 15 | 5jk2-A | 7.1 | 2.4 | 85 | 128 | 9 | TP0751 |
| 16 | 4rlc-A | 7.1 | 3.9 | 108 | 135 | 6 | OUTER MEMBRANE PORIN F |
| 17 | 2m4l-A | 7 | 3.1 | 84 | 99 | 8 | PROTEIN BT_0846 |
| 18 | 4pr7-A | 6.9 | 3.4 | 89 | 171 | 11 | OLIGOGALACTURONATE-SPECIFIC PORIN KDGM |
| 19 | 2jw1-A | 6.8 | 3 | 91 | 115 | 12 | LIPOPROTEIN MXIM |
| 20 | 5ha0-A | 6.4 | 2.4 | 89 | 156 | 7 | LIPOCALIN AI-4 |

